# Identification of the host reservoir of SARS-CoV-2 and determining when it spilled over into humans

**DOI:** 10.1101/2023.11.25.568670

**Authors:** Vidyavathi Pamjula, Norval J.C Strachan, Francisco J. Perez-Reche

**Affiliations:** The School of Natural and Computing Sciences - Department of Physics, University of Aberdeen, Scotland, UK

**Author notes:** Corresponding author. Vidyavathi Pamjula, PhD student.

## Abstract

Since the emergence of SARS-CoV-2 in Wuhan in 2019 its host reservoir has not been established. Phylogenetic analysis was performed on whole genome sequences (WGS) of 71 coronaviruses and a Breda virus. A subset comprising two SARS-CoV-2 Wuhan viruses and 8 of the most closely related coronavirus sequences were used for host reservoir analysis using Bayesian Evolutionary Analysis Sampling Trees (BEAST). Within these genomes, 20 core genome fragments were combined into 2 groups each with similar clock rates (5.9×10^−3^ and 1.1×10^−3^ subs/site/year). Pooling the results from these fragment groups yielded a most recent common ancestor (MRCA) shared between SARS-COV-2 and the bat isolate RaTG13 around 2007 (95% HPD: 2003, 2011). Further, the host of the MRCA was most likely a bat (probability 0.64 - 0.87). Hence, the spillover into humans must have occurred at some point between 2007 and 2019 and bats may have been the most likely host reservoir.

## 2 Introduction

Emerging and re-emerging infectious diseases caused by viruses (e.g. severe acute respiratory syndrome (SARS), Middle East respiratory syndrome (MERS) ^1^, Influenza ^2,3^), bacteria (e.g. Lyme disease ^4^, Cholera ^5^, Plague ^6^, *Escherichia coli* O157:H7 (E. coli)^7–9)^, fungi (e.g. *Cryptococcus gatti* infection ^10^) and parasites (e.g. malaria ^11^) are still the leading causes of death globally ^12^. They also raise concerns about global health, biosecurity and economic disruption ^13,14^. Zoonoses comprise 60% of total infectious diseases whose agent is transmitted from animal hosts to humans ^15^.

Coronaviruses belong to the sub-family Coronavirinae and can be zoonotic ^16–18^. They have caused two major human epidemics in the two decades preceding the SARS-CoV-2 pandemic: severe acute respiratory syndrome (SARS) in 2002-2003 and Middle East respiratory syndrome (MERS) in 2012. Research indicates SARS-CoV are closely related to viruses isolated from Chinese horseshoe bats (*Rhinolophus sp.)* which are considered to be the likely reservoir host ^19–21^. *Vespertilio superans* bats are also thought to be the primary reservoir for MERS ^22^. Intermediate hosts have been postulated for both of these epidemics. For example, coronavirus isolates from Himalayan palm civet cats were highly homologous to human SARS-CoV ^17,23,24^ as were dromedary camels for MERS-CoV ^25–27^. However, whether civet-cats/dromedary camels were actually intermediate hosts remains an open question.

A human outbreak of Novel Coronavirus (SARS-CoV-2) with cases presenting symptoms of pneumonia was reported on December 8^th^ 2019 in Wuhan, China. These cases were epidemiologically associated with a fresh seafood and wild animal market in Wuhan ^28–30^. By January 7^th^ 2020, the agent of this pneumonia was isolated from the respiratory epithelium of patients ^31,32^ and on 11^th^ February WHO named this new coronavirus as severe acute respiratory syndrome coronavirus 2 (SARS-CoV-2) ^33,34^. Subsequently, there was worldwide human transmission resulting in a global pandemic ^35^ and as of 4^th^ October 2023, there has been 771,151,224 infections and 6,960,783 deaths reported ^36^. The International Monetary Fund estimates cumulative output loss from the pandemic through to 2024 at $13.8 trillion ^37^.

Coronaviruses comprise four genera: alpha, beta, gamma and delta. Alpha and Betacoronaviruses are frequently found in bats and are mainly associated with infections in mammals ^38^. The Betacoronavirus genus includes SARS-CoV-2, SARS CoV and MERS ^34^ which can cause respiratory, gastrointestinal, hepatic and central nervous system infections in humans ^39^. Host species for the Gammacoronavirus genus include birds and Beluga whales. Deltacoronaviruses have been found in birds and mammals ^40,41^.

Studies utilizing whole genome sequencing have enabled an understanding of the phylogenies, transmission, genetic diversity and outbreak dynamics of zoonotic viruses including: SARS CoV ^14,39^; MERS CoV ^18,42^; Ebola ^43^; HIV ^44^ and influenza ^45,46^. These studies utilised Bayesian phylodynamic methods ^47^ to determine molecular clock rate(s) and likely host reservoirs as well as elucidating the temporal and geographical patterns of transmission.

Identification of the temporal signal between genome sequences is crucial to obtain timed phylogenies. The analysis of SARS-CoV-2 has proven challenging in this respect and no evidence of a temporal signal was obtained in previous studies ^48^. This might have affected the accuracy of former estimates for the time to the most recent common ancestor. The lack of a temporal signal may be due to recombination and/or different molecular clock rates in different parts of the genome. To circumvent this problem, new methods need to be developed to identify genome fragments that exhibit a similar molecular clock rate across coronavirus genomes.

Bats have been postulated as a likely reservoir from which SARS-CoV-2 originated. Evidence pointing in this direction includes the close genetic relationship between a coronavirus isolated from *Rhinolophu*s bats in Yunnan, Southern China ^49^ and SARS-CoV ^50^. At the whole genome sequence level, the closest available sequence to SARS-CoV-2 to date is the RaTG13 virus sampled from a *Rhinolophus affinis* bat (96.2% similarity ^32,51^). Also, serological surveillance of people living in villages close to the natural habitat of bats in caves revealed 2.9% bat coronavirus seroprevalence in humans. This indicates that human exposure to bat-CoVs may be relatively common in China and that there is an opportunity for these viruses to spill over directly into humans without the need for an intermediate host ^50^. However, there is the possibility of an intermediate host reservoir between bats and humans^51^. Several have been hypothesised for SARS-CoV-2. These include rodents ^52^, racoon dogs ^53^, pangolin ^54,55^ and other animals ^56^. Despite significant efforts to identify an intermediate host, none has been clearly identified so far.

The zoonotic reservoir of SARS-CoV-2, the length of time the SARS-CoV-2 lineage has circulated in the host reservoir and the time when the first transmission occurred into humans remain unclear. This paper aims to: (1) identify the host reservoir of SARS-CoV-2, (2) to determine when SARS-CoV-2 spilled over from the host reservoir to humans and (3) to determine the phylogenetic relationship between SARS-CoV-2 and other coronaviruses. This will be achieved by conducting phylogenetic and Bayesian phylodynamic analysis utilizing a database of 71 coronavirus genomes and identification of core genome fragments with indistinguishable molecular clock rates.

## 3 Methods

### Data

Coronavirus whole genome sequences (n = 71) and a Breda virus genome sequence were obtained from Genbank, the National Centre for Biotechnology Information (NCBI) and the Global Initiative on Sharing All Influenza Data (GISAID) for phylogenetic analysis (see Table S1). Metadata including year of isolation, host and country were also collected. Three sub-samples (selections) of these sequences were used to address the aims of this paper (Figure 1).

**Figure 1:**
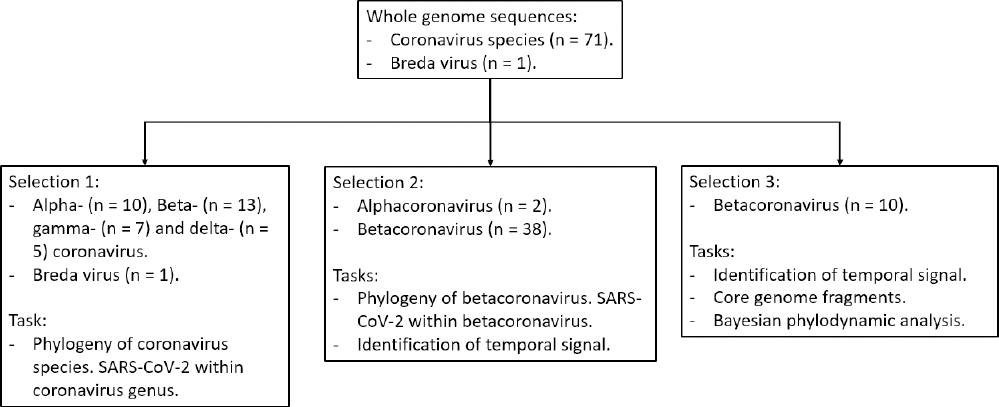
Genomes used in this study. Three selections of the 72 genomes in the original dataset were used for phylogenetic and phylodynamic analyses. Details on the genomes within each selection are given in Table S1.

### 3.1 Phylogeny of coronavirus

A selection of 35 whole genome sequences representative of the Alpha, Beta, Gamma and Delta coronavirus genera and a Breda virus genome were used to determine the location of SARS-CoV-2 Wuhan viruses within the coronavirus genus (Table S1, Selection 1).

A subset of 40 sequences (Table S1, Selection 2) was used to determine the potential origin of the outbreak of SARS-CoV-2. These sequences comprise Betacoronaviruses (n=38, SARS-CoV-2, SARS-CoV, MERS-CoV, and viruses from a wide range of hosts) along with Alphacoronaviruses (n=2) as the outgroup.

Genome selections 1 and 2 were aligned using ClustalW ^57,58^.The alignment was used to generate phylogenies with MEGA X^59^ using the neighbour-joining method with bootstrapping (1000 replications).

### 3.2 Identification of temporal signal from coronavirus genomic data

#### 3.2.1 Whole Genome Sequences

40 WGS (Table S1, Selection 2) and 10 WGS (Table S1, Selection 3) were used to assess the temporal signal utilising root-to-tip regression by TempEST^60^.

#### 3.2.2 Core genome fragments

##### 3.2.2.1 Identification of core genome fragments

Identification of core genome fragments was performed on a subset of the coronavirus genome sequences (n=10) (Table S1, Selection 3). Multiple sequence alignments for the 10 genomes were again obtained using ClustalW. Contiguous fragments with more than 200 consecutive nucleotide base positions (bp) across the 10 WGS were identified (Figure 2(a)). The identified core genome fragments were mapped to the reference sequence of SARS-CoV-2 (Gen Bank -NC-045512.2) to determine the non-structural proteins present.

##### 3.2.2.2 Identification of core genome fragment groups with indistinguishable molecular clock rates

###### 3.2.2.2.1 Pairs of fragments

The pairwise genetic distance between sequences *i* and *j* in a fragment *f* was measured by the number 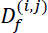 of nucleotide differences between them. Figure 2(b) shows the pairwise genetic distance, 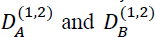 between sequence 1 and sequence 2, for fragments *A* and *B* respectively.

The pairwise genetic distance 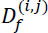 was normalized to account for the fact that every fragment identified across 10 WGS had a different number of nucleotides. More explicitly, the pairwise distance between two sequences *i* and *j* corresponding to a fragment *f* was estimated by the proportion of sites that differ between the two sequences in this fragment, i.e.

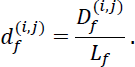

Here, *L_f_* ≥ 200 is the length of fragment *f*.

**Figure 2:**
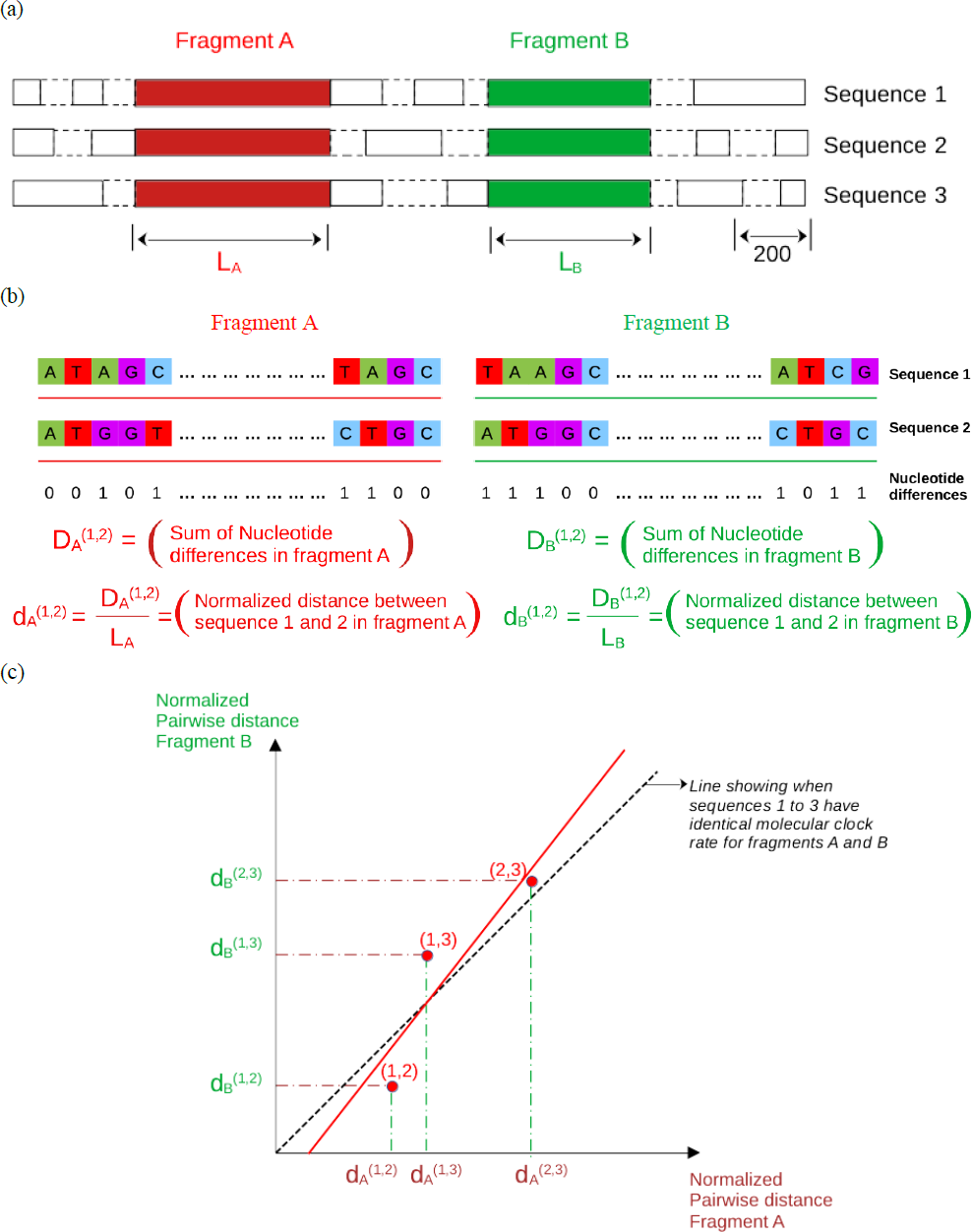
Method to identify genome fragment groups with indistinguishable molecular clock rates exemplified for three whole genome sequences. (a) Identification of contiguous DNA core fragments with more than 200 consecutive nucleotide base positions (rectangles with solid border indicate nucleotide fragments; dashed lines indicate genome gaps). This example identifies two fragments (*A* and *B*) from the three genomes. (b) Calculation of normalised pairwise genetic distance 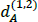 and 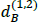 between sequence 1 and sequence 2 in terms of fragment *A* and fragment *B* respectively. (c) Virtual example of linear regression analysis using normalized pairwise distances between fragments *A* and *B* across the three sequences. The dashed line indicates the ideal case in which fragments *A* and *B* have identical molecular clock rates.

###### 3.2.2.2.2 Identifying groups of fragments with indistinguishable clock rates

The normalized pairwise distance between pairs of fragments was used to identify groups of fragments with statistically indistinguishable molecular clock rates. In a hypothetical scenario in which two fragments *A* and *B* follow the same molecular clock for a pair of sequences *i* and *j*, it would be expected that 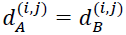. If all the sequences evolve with the same molecular clock according to fragment *A* and *B*, the distances 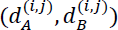 between any pair of sequences (*i*, *j*) should fall along the line with zero intercept and slope 1 in the space (*d_A_*,*d_B)_*. In general, the points 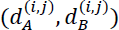 for all pairs of sequences will not exactly fall along the ideal straight line (see an example in Figure 2 (c)). In practice, a line was fitted to the points 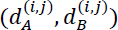 for all pairs of sequences and the fragments *A* and *B* were assumed to follow the same molecular clock if the fitted line is close enough to the ideal line. More specifically, the fitted line was considered to agree with the ideal line if (i) the slope was statistically consistent with 1, (ii) the intercept was statistically compatible with 0 and (iii) the p-value for the correlation coefficient was smaller than 0.05. Only pairs of fragments satisfying these conditions were assumed to follow indistinguishable molecular clock rates. From a statistical viewpoint, these pairs of fragments are considered evolutionary synchronous. The statistical significance of the hypotheses slope = 1 and intercept = 0 were tested in terms of a 95% confidence interval. The fragment groups identified by this method were used in the BEAST analysis described below.

### 3.3 Bayesian phylodynamic analysis by BEAST

#### 3.3.1 Bayesian phylodynamic analysis to estimate clock rate and timed phylogeny

Datasets comprising 10 WGS, all core genome fragments and the fragment groups with indistinguishable molecular clocks were used to conduct Bayesian phylodynamic analyses with BEAST (v.2.6.2)^47,61^. This involved three sub-models. The First is the DNA substitution sub-model which has 2 options: (1) Hasegawa-Kishino-Yano (HKY) and (2) General Time Reversible (GTR) model. The second is the population sub-model which has 4 options: (1) Coalescent Constant Population (CCP); (2) Coalescent Exponential Population (CEP); (3) Coalescent Bayesian Skyline (CBS) and (4) Yule Model. The third is the molecular clock sub-model with 2 options: (1) strict (SC) and (2) relaxed (RC) molecular clock. Both SC and RC had uninformed prior distributions. The clock rates obtained from this analysis were used to specify the clock rate priors in the subsequent host probability calculations ^62–64^ of section 3.3.3.

Models were run using the Markov Chain Monte Carlo (MCMC) method with 10^8^ iterations after 10% burn-in and sampled once every 1000 iterations. All permutations of substitution, population and clock models (n=16) for each dataset (n=4) were performed. Hence, in total 64 runs were carried out in the BEAST analysis. Convergence was assessed using the effective sample size (ESS) criterion (ESS>100) implemented on Tracer v.1.71^65^. Tree Annotator was used to produce a Maximum Clade Credibility (MCC) tree and from this tMRCA of SARS-CoV-2 Wuhan, with 95% HPD was obtained. The timed phylogeny was visualized using Fig Tree v 1.4.4^66,67^.

#### 3.3.2 Model selection using Nested sampling

Nested sampling (NS) was used to identify the best fitting parsimonious models^68,69^. This method uses two tuning parameters: The number of points sampled from the prior (number of particles, set to n=32) and chain length set to 20,000 steps with sub-chain length of 10,000 steps. Only model runs that converged in the previous section were used in this analysis.

#### 3.3.3 Determination of host reservoir probability, molecular clock rate and time to most recent common ancestor

To estimate the most probable host reservoir of SARS-CoV-2 and the time period of spill over from animal reservoirs to humans, host transition and evolutionary dynamics was performed using BEAST (v1.10.4) which implements a method to estimate cross-species transmission ^70^. Models selected by NS together with priors using the clock rate obtained as described above were used in the analysis. Posterior distributions were obtained through MCMC analysis runs with chain lengths of 1 x 10^8^ generations and convergence was assumed to be achieved when ESS >100. If ESS >100 was not achieved, the MCMC chain length was increased to 2 x 10^8^ generations and the ESS was checked.

##### 3.3.3.1 Determination of pooled estimate of tMRCA and associated uncertainty

All the estimates for the time to most recent common ancestor (tMRCA) and their associated uncertainties, obtained by the selected models with NS, were pooled using propagation of errors ^71^.

## 4 Results and discussion

### 4.1 Phylogenetic analysis

#### 4.1.1 Classification of SARS-CoV-2 within Coronavirus

The phylogenetic analysis of coronavirus showed that the four different genera of coronavirus (alpha, beta, gamma and delta) belong to four distinct clades (Figure 3(a)). SARS-CoV-2 belongs to the Betacoronavirus clade as reported previously ^34^.

#### 4.1.2 Phylogenetic relationship between SARS-CoV-2 Wuhan and other beta coronaviruses

Phylogenetic analysis (Figure 3 (b)) showed that all SARS-CoV-2 Wuhan genomes grouped together and that Bat CoV RaTG13 (obtained from bat *Rhinolophus affinis*) ^23^ was the most closely genetically related to SARS-CoV-2. The next most closely related to SARS-CoV-2 was a virus sampled from Pangolin (*Manis javanica* – MP789)^72^. This agrees with previous research which found that RaTG13 ^48,54^ was the most closely related virus to SARS-CoV-2 with 96.2% whole genome homology ^32,51^. These authors also found Pangolin-CoV as the next most closely related with 91.02% WGS similarity^54^. SARS-CoV are the next closest group to the SARS-CoV-2 clade with 79.5% WGS similarity, see Figure 3(b) ^32,73^.

MERS-CoV is phylogenetically more distant from the SARS-CoV-2 Wuhan clade than SARS-CoV and only shares 50% sequence identity, see Figure 3(b)^73,74^. The Alphacoronavirus samples are clustered together at the root of the tree as an outgroup, as expected.

**Figure 3:**
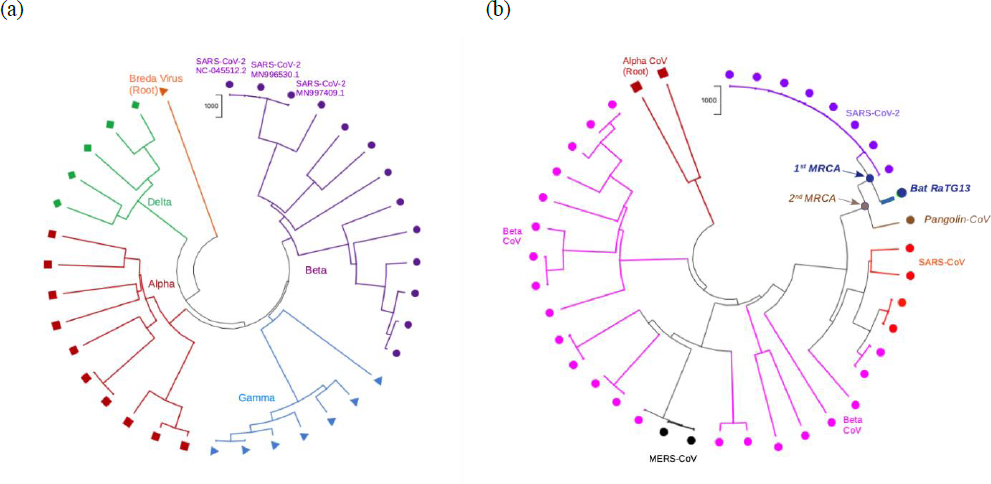
Phylogenetic analyses. (a) Neighbour-joining tree of 36 WGS representative of coronavirus (Alpha – maroon; Beta – purple; Gamma – blue and Delta – green). The tree is rooted with Breda virus (peach). Three genome sequences of SARS-CoV-2 (purple) cluster within the Betacoronavirus clade (see Table S1 - Data Selection 1) (b) Neighbour-joining tree of Betacoronavirus (n=38) with Alphacoronavirus samples (n=2, maroon) as root (see Table S1 - Data Selection 2); SARS-CoV-2 (purple); SARS-CoV (red), MERS-CoV (black), other Betacoronaviruses (Beta CoV, pink). BatCoV RaTG13 (indigo) is the 1^st^ MRCA and Pangolin coronavirus (brown) is the 2^nd^ MRCA to SARS-CoV-2 (purple). The scale bars provide 1000 bootstraps in the MEGA X.

### 4.2 Identification of temporal signal from coronavirus genomic data

#### 4.2.1 Whole Genome Sequences

Temporal analysis using TempEst utilising 40 WGS and 10 WGS (see Table S1 - Selections 2 and 3, respectively) failed to find a statistically significant correlation coefficient in terms of the root-to-tip divergence (Supplementary Figure 1, Table S2). This suggests a lack of temporal signal for both datasets, in agreement with other studies ^48^. This may be due to the fact that these viruses are from a diverse coronavirus population with deep evolutionary histories.

#### 4.2.2 Core genome fragments

##### 4.2.2.1 Identification of core genome fragments

Twenty fragments with *L_f_* ≥ 200 were identified from the 10 WGS alignment of 32794 base pairs (bp) (Figure 4). The genomic location of the identified 20 fragments was found to be within a 266 to 21555 bp region which corresponds to two open reading frames 1ab (ORF’s)^75–77^. The ORF1ab gene encodes non-structural proteins (nsps): nsp 1 to 11 and nsp 12 to 16, respectively (Table S3).

**Figure 4:**
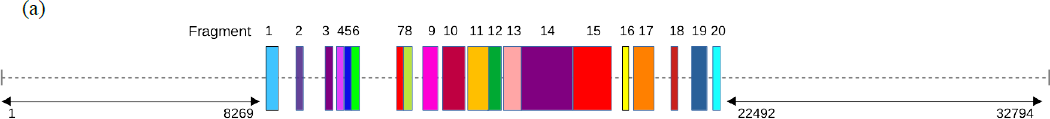
Scale diagram of the core genome fragments (n=20) with more than 200bp from the 10 WGS multiple sequence alignment (Tables S1, Selection 3 and S3).

##### 4.2.2.2 Identification of core genome fragment groups with indistinguishable molecular clock rates

The 20 core genome fragments obtained above were analysed to form groups of fragments with indistinguishable molecular clock rates using the criteria in Section 3.2.2.2.2. Two groups of three fragments were identified with a mutually consistent temporal signal: {1,5,16} and {8,11,12} (Table 1). Scatterplots for the normalised pairwise distances for these fragments are presented in Supplementary Figures 2 and 3.

**Table 1:** Regression analysis parameters for core genome fragment groups which satisfy the conditions of indistinguishable molecular clock rate.

| Fragment Group | Fragment number | Fragment number | p-value | Slope | Slope (95% CI) | Intercept | Intercept (95% CI) |
| --- | --- | --- | --- | --- | --- | --- | --- |
| 1 | Frag 1 | Frag 5 | 1.64E-16 | 0.89 | (0.75, 1.02) | 1.25 | (-1.18, 3.69) |
|  | Frag 1 | Frag 16 | 3.74E-24 | 0.92 | (0.83, 1.01) | 0.35 | (-1.23, 1.93) |
|  | Frag 5 | Frag 16 | 3.46E-19 | 0.90 | (0.78, 1.01) | 1.25 | (-0.77, 3.26) |
| 2 | Frag 8 | Frag 11 | 9.55E-33 | 0.98 | (0.92, 1.03) | 0.04 | (-0.76, 0.84) |
|  | Frag 8 | Frag 12 | 4.13E-25 | 0.94 | (0.86, 1.03) | -0.43 | (-1.62, 0.77) |
|  | Frag 11 | Frag 12 | 1.52E-31 | 0.97 | (0.91, 1.03) | -0.50 | (-1.34, 0.33) |

### 4.3 Bayesian Phylodynamic analysis by BEAST

#### 4.3.1 Model selection using Nested Sampling

Nested sampling found that the HKY substitution model provided the best fits for all the datasets. In contrast, the best combination of population and molecular clock models was found to be data specific. This resulted in six evolutionary models being selected (Table 2). For the 10 WGS dataset, the best fitting model was one with coalescent exponential population and relaxed molecular clock (CEP-RC). Nested sampling was unable to discriminate between the fits of two combinations of models when using the 20 core fragments dataset: Coalescent constant population with relaxed clock (CCP-RC) and coalescent exponential population with relaxed clock (CEP-RC). Both model combinations are taken as the best fits for this dataset. The best fit to the evolution of the fragment group {8,11,12} was also obtained for two combinations of models which are statistically indistinguishable: the coalescent population model combined with a strict clock model (CCP-SC) and a coalescent constant population model combined with a relaxed clock model (CCP-RC). The combination CCP-SC also provided the best fit to the evolution of the fragment group {1,5,16}. By estimating maximum Effective Sample Size (Max ESS) using Nested sampling, increasing the number of particles to 32 in the parameter space at each iteration, yields more precision to reach better convergence. This provides higher values of Max ESS estimate for six evolutionary models (Supplementary Figure 4).

**Table 2.** Results of the Bayesian phylodynamic and host probability analyses for the selected 6 best fitting models identified by nested sampling. The final row provides the pooled values from the 6 models.

| Data and evolutionary model | Molecular clock rate [95% CI] | Year of 1st MRCA [95% CI] | 1st MRCA host probability (human, pangolin, bat) | Year of 2nd MRCA [95% CI] | 2nd MRCA host probability (human, pangolin, bat) | ESS |
| --- | --- | --- | --- | --- | --- | --- |
| 10WGS, CEP-RC | 0.0029<br>[0.0018, 0.0049] | 2006<br>[2000, 2011] | (0.16, 0.02, 0.82) | 1998<br>[1987, 2007] | (0.14, 0.08, 0.79) | 445 |
| 20 FG, CCP-RC | 0.0039<br>[0.0022, 0.0079] | 2009<br>[2002, 2013] | (0.13, 0.01, 0.86) | 2003<br>[1985, 2011] | (0.12, 0.07, 0.81) | 124 |
| 20 FG, CEP-RC | 0.0059<br>[0.0041, 0.0095] | 2010<br>[2005, 2013] | (0.11, 0.02, 0.87) | 2005<br>[1996, 2011] | (0.10, 0.08, 0.82) | 818 |
| {8,11,12} FG, CCP-RC | 0.0058<br>[0.0032, 0.0116] | 2010<br>[2004, 2013] | (0.14, 0.02, 0.84) | 2006<br>[1995, 2012] | (0.13, 0.06, 0.81) | 1651 |
| {1,5,16} FG, CCP-SC | 0.0023<br>[0.0020, 0.0030] | 2005<br>[2000, 2009] | (0.28, 0.01, 0.70) | 1984<br>[1973, 1996] | (0.26, 0.16, 0.58) | 62571 |
| {8,11,12} FG, CCP-SC | 0.0011<br>[0.0010, 0.0014] | 1998<br>[1992, 2004] | (0.34, 0.02, 0.64) | 1972<br>[1959, 1985] | (0.29, 0.18, 0.52) | 62671 |
| Pooled values | 0.0037<br>[0.0017, 0.0057] | 2007<br>[2003, 2011] | (0.19 [0.16, 0.23],<br>0.02 [0.02, 0.02],<br>0.79 [0.75, 0.83]) | 1995<br>[1986, 2004] | (0.17 [0.14, 0.21],<br>0.11 [0.08, 0.13],<br>0.72 [0.67, 0.78]) | 21380 |

#### 4.3.2 Clock rate, time to the most recent common ancestor and host reservoir probability

##### Clock rate

Clock rates for the six selected evolutionary models range from 1.1 to 5.9× 10^−3^ subs/site/year (Table 2). The clock rate of the two models using a strict clock ({1,5,16}-CCP-SC and {8,11,12}-CCP-SC) tends to be slower on average than that of models with a relaxed clock. However, only the model based on the fragment group {8,11,12} is statistically slower than the rest (Supplementary Figure 5(a)). Pooling the values of the molecular clock rate from the six models yielded 0.0037 (0.0017 - 0.0057) subs/site/year. This estimate is compatible with the molecular clock rate of SARS, 0.00169 (0.00131 - 0.00205) subs/site/year found previously by Boni et al^48^. In the same work, however, the authors used a slower estimate of the molecular clock rate, 0.00055 subs/site/year, for their phylogenetic dating methods. This estimate was obtained for non-recombinant regions of a 68-genome sarbecovirus alignment using the molecular clock rate of MERS-CoV (0.00078 subs/site/year) and HCoV-OC43 (0.00024 subs/site/year) to define a prior. Such estimates were used to circumvent the lack of temporal signal in their dataset. There are several reasons that could explain the difference between the current studies pooled estimate of the molecular clock rate and the estimate obtained by Boni et al.,^48^. These include the selection of genomes and the parts of the genome used. Another possibility is that the estimated molecular clock rate is influenced by the relatively slow clock rate of the prior based on MERS-CoV and HCoV-OC43. In the current study, by estimating the clock rate distribution for each of the six datasets (Table S5) yields a faster pooled estimate of molecular clock rate.

##### Time to the most recent common ancestor

All of the 6 models selected by nested sampling gave statistically compatible estimates for the time to the 1^st^ MRCA of SARS-CoV-2 and BatCoV RaTG13 (Table 2 and Supplementary Figure 5(b)). Pooling the results from the six models (Table S4(a)), the time to the 1^st^ MRCA was found to be 13.5 ± 4.1 years which corresponds to the year 2007 (2003 - 2011) (Supplementary Figure 6). This estimate for the time to the 1^st^ MRCA is more recent than the estimates given in Boni et al.,^48^ : 1948 (1879 – 1999), 1969 (1930 – 2000) and 1982 (1948 – 2009). This is expected due to the slower molecular clock rate used, as discussed above.

The time to the 2^nd^ MRCA between Pangolin-CoV and the SARS-CoV-2 Wuhan/bat RaTG13 lineage was also statistically similar across the six models (Supplementary Figure 5(b)). Pooled results from the six models estimated it to be 25.4 ± 8.9 years, i.e., [1995 (1986 - 2004)], (Table 2, Table S4(b)). This estimate is also more recent than 1851(1730 - 1958) determined by Boni et al.,^48^ .

##### Host reservoir probability

The most likely host reservoir of the 1^st^ MRCA between SARS-CoV-2 and BatCoV RaTG13 was a bat with a probability of 0.79 [0.75, 0.83] (pooled over the six selected models, Table 2). This indicates bat as potentially the natural zoonotic origin of SARS-CoV-2. The pooled probability of origin for humans was 0.19 [0.16, 0.23] and pangolin 0.017 [0.015, 0.019] (Supplementary Figures 7 and 8). From the estimated time for the 1^st^ MRCA and the likelihood of bat as a natural reservoir, it can be concluded that the ancestor of SARS-CoV-2 spilled over from bats to humans sometime between 2007 and 2019. It is unclear whether that was directly into humans or via an intermediate animal reservoir.

##### Evidence of bats as the natural reservoir

The results (Table 2) agree with the finding that phylogenetically close coronaviruses to SARS-CoV-2 have been circulating in bats for many decades^48,78^. Despite the likelihood of a spill-over of SARS-CoV-2 from the bat reservoir to humans, there is a lack of intermediate sequences that prevents a precise understanding of the spill over in terms of whether a potential intermediate host exists (WHO, 2021). The first recorded human cases with severe pneumonia were at Jin Yin-Tan hospital in Wuhan in December, 2019 and full genomes were sequenced at Wuhan Institute of Virology (WIV) (Zhou et al., 2020). However, the high prevalence of asymptomatic and unreported cases^79^ and the low surveillance prior to the Wuhan outbreak make the identification of an intermediate host, even if one existed, very challenging.

## 5 Conclusion

Coronaviruses closely related to SARS-CoV-2 appear to have emerged from bats at some point within the 13 years before the SARS-CoV-2 outbreak. This is more recent than the estimates from previous studies which suggested 40 to 70 years ^48^. Bat was found to be the most likely natural reservoir for SARS-CoV-2. There may be an intermediate host reservoir between bats and humans but there is insufficient information to demonstrate this hypothesis. To be more precise about the spillover of SARS-CoV-2 into humans, more sequencing and epidemiological evidence would be required. Unfortunately, this is difficult to achieve retrospectively and would have required an internationally coordinated surveillance program prior to the pandemic. To prevent future zoonotic pandemics, such a surveillance system will be required. This will require collaboration between epidemiologists, geneticists, medics, mathematical modelers and social scientists.

## Supporting information

R Program

10 WGS Dataset

## Acknowledgements

FJPR acknowledges funding from a Medical Research Council Fellowship (MR/W021455/1).

For the purpose of open access, the author has applied a Creative Commons Attribution (CC BY) licence to any Author Accepted Manuscript version arising from this submission.

## Author contributions

VP contributed to the ideas behind this work, carried out the analysis and led the writing of the paper. FJPR and NJCS contributed to the ideas behind the work, advised on the analysis and edited drafts of the paper.

