## Supplementary Information for "Identification of the host reservoir of SARS-CoV-2 and determining when it spilled over into humans"

***Supplementary Tables:***

Table S1: Metadata of Coronavirus Genomes 2

Table S2: Temporal exploration of 40 and 10 whole genome sequences using TempEST 3

Table S3: Characteristics of 20 core genomic fragments of length $Lf \geq 200$ obtained from 10 WGS. Number and percentage of Single Nucleotide Polymorphism (SNP's). Non-structural proteins (NSP's) identified from the 20 fragments along with their positional mapping to reference sequence of SARS-CoV-2 NC_045512.2 is presented. 4

Table S4: 4a, 4b and 4c - Propagation of Errors for 1$^{st}$ tMRCA, 2$^{nd}$ tMRCA and clock rate 5

Table S5: Bayesian analysis clock rate estimates 6

***Supplementary Figures:***

Supplementary Figure 1: Temporal Exploration of Sequences and Trees [TempEST] performed on Whole Genome Sequences 7

Supplementary Figure 2: Regression analysis of the normalized pairwise distance for fragment group {1,5,16} 8

Supplementary Figure 3: Regression analysis of the normalized pairwise distance for fragment group {8, 11, 12**}** 9

Supplementary Figure 4: BEAST Model selection by Nested Sampling - Estimation of Max effective sample size (ESS) 10

Supplementary Figure 5: BEAST Phylodynamic evolutionary analysis for the evolutionary models selected by Nested sampling 11

Supplementary Figure 6: BEAST Phylotraits Evolutionary Analysis maximum clade credibility phylodynamic trees 14

Supplementary Figure 7: BEAST Phylotraits evolutionary analysis – Host probability for SARS-CoV-2 and Pangolin CoV 15

Supplementary Figure 8: BEAST Phylotraits evolutionary analysis Host Probability of MRCA to SARS-CoV-2 16

### Supplementary Tables

#### Table S1: Metadata of Coronavirus Genomes

Listed below are the genomes used in the study: 71 coronavirus genomes and a Breda virus genome. Selection 1, comprising 36 sequences was used to generate a phylogeny across all the species (alpha, beta, gamma and delta) of coronavirus. Selection 2 comprising 40 sequences was used to generate a phylogeny across the beta coronaviruses. Selection 3, comprising 10 beta coronavirus sequences closest to SARS-CoV-2 was used in the BEAST phylodynamic analysis.

| Seq. No | Dataset Selection | | | Group | Source | Country | Collection/ Submission Year | GenBank/ GISAID Sequence |
|---|---|---|---|---|---|---|---|---|
| | 1 | 2 | 3 | | | | | |
| 1 | X | | | Gamma | Beluga Whale coronavirus SW1 | USA | 2007 | NC_010646.1 |
| 2 | X | | | Gamma | Infectious Bronchitis Avian coronavirus | USA | 2004 | MN711790.1 |
| 3 | X | | | Gamma | Avian infectious Bronchitis coronavirus | UK | 2004 | NC_ 001451.1 |
| 4 | X | | | Gamma | Turkey coronavirus TCoV-ATCC | USA | 2007 | EU022526.1 |
| 5 | X | | | Gamma | Turkey coronavirus TCoV/TX-GLO/01 | USA | 2001 | GQ427174.1 |
| 6 | X | | | Gamma | Turkey coronavirus TCoV/TX1038/98 | USA | 1998 | GQ427176.1 |
| 7 | X | | | Gamma | Turkey coronavirus TCoV/TX1038/98/2 | USA | 1998 | GQ427176.1 |
| 8 | X | | | Delta | Bulbul coronavirus HKU 11-934 | Hong Kong | 2007 | NC_ 011547.1 |
| 9 | X | | | Delta | Thrush coronavirus HKU12-600 | Hong Kong | 2007 | NC_011549.1 |
| 10 | X | | | Delta | Munia coronavirus HKU13-3514 | Hong Kong | 2007 | NC_011550.1 |
| 11 | X | | | Delta | Coronavirus HKU15 | Hong Kong | 2014 | LC216915.1 |
| 12 | X | | | Delta | Sparrow coronavirus HKU17 | Hong Kong | 2007 | NC_016992.1 |
| 13 | X | | | Alpha | Porcine respiratory – PRCV ISU-1 | USA | 2006 | DQ811787.1 |
| 14 | X | | | Alpha | Pig – Transmissible gastroenteritis virus | USA | 2001 | NC _038861.1 |
| 15 | X | | | Alpha | Feline infectious peritonitis virus | USA | 2005 | NC_ 002306.3 |
| 16 | X | | | Alpha | Human coronavirus 229E | Germany | 2000 | NC_ 002645.1 |
| 17 | X | | | Alpha | Human coronavirus 229E | Germany | 2001 | AF304460.1 |
| 18 | X | | | Alpha | Human coronavirus NL63 | Netherlands | 2004 | NC_ 005831.2 |
| 19 | X | | | Alpha | Bat coronavirus HKU-8 | Hong Kong | 2008 | NC _010438.1 |
| 20 | X | | | Alpha | Porcine epidemic diarrhea virus | Italy | 2016 | KY111278.1 |
| 21 | X | | | Alpha | Scotophilus bat coronavirus 512 | Hong Kong | 2005 | NC_009657.1 |
| 22 | X | | | Breda | Breda virus - Torovirus | Canada | 2003 | NC_007447.1 |
| 23 | X | | | Beta | Mutant Bat SARS Coronavirus HKU3 | USA | 2008 | MN782115.1 |
| 24 | X | | | Beta | Rousettus Bat Coronavirus HKU9 | China | 2009 | MG762674.1 |
| 25 | X | | | Beta | Human coronavirus HKU1 | USA | 2010 | KF686346.1 |
| 26 | X | | | Beta | Murine coronavirus MHV-JHM.1A | USA | 2009 | FJ647226.1 |
| 27 | X | | | Beta | Equine coronavirus | Japan | 2012 | LC061274.1 |
| 28 | X | | | Beta | Human coronavirus HCoV-OC43 | UK | 2011 | KU131570.1 |
| 29 | X | | | Beta | Bovine coronavirus BCoV-2014_13 | France | 2014 | KX982264.1 |
| 30 | X | | | Beta | Bovine coronavirus BCoV-ENT | USA | 2001 | AF391541.1 |
| 31 | X | | | Beta | Homo sapiens - SARS-CoV-2 WIV06 | China | 2019 | MN996530.1 |
| 32 | X | | | Beta | Homo sapiens – SARS-CoV-2 | USA | 2020 | MN997409.1 |
| 33 | X | X | | Alpha | Rhinolophus bat coronavirus HKU2 | China | 2006 | EF203064.1 |
| 34 | X | X | | Beta | Tylonycteris bat coronavirus HKU41 | China | 2006 | EF065505.1 |
| 35 | X | X | | Beta | Pipistrellus bat coronavirus HKU5 | China | 2006 | EF065509.1 |
| **36*** | **X** | **X** | **X** | **Beta *** | **Homo sapiens SARS-CoV-2 Wuhan-Hu1*** | **Wuhan - China** | **2020** | **NC_045512.2 *** |
| 37 | | X | X | Beta | Homo sapiens SARS | Canada | 2003 | NC_004718.3 |
| 38 | | X | X | Beta | Bat/ Yunan/ RaTG/2013/EPI_ISL 402131 | China | 2013 | EPI_ISL_402131 |
| 39 | | X | X | Beta | Homo sapiens SARS-CoV-2 Wuhan-Hu-1 | Wuhan - China | 2019 | MN908947.3 |
| 40 | | X | X | Beta | Bat SARS-like coronavirus RsSHC014 | China | 2011 | KC881005.1 |
| 41 | | X | X | Beta | Homo sapiens -SARS coronavirus BJ01 | China | 2003 | AY278488.2 |
| 42 | | X | X | Beta | Bat coronavirus BM48-31/BGR/2008 | Bulgaria | 2008 | NC_014470.1 |
| 43 | | X | X | Beta | Bat SARS coronavirus Rf1 | China | 2004 | DQ412042.1 |
| 44 | | X | X | Beta | Bat SARS-like coronavirus WIV1 | China | 2012 | KF367457.1 |
| 45 | | X | X | Beta | Pangolin coronavirus MP789 | China | 2019 | MT084071.1 |
| 46 | | X | | Beta | Human Middle East respiratory syndrome | Middle East | 2012 | JX869059.2 |
| 47 | | X | | Beta | Human coronavirus HKU1 | China | 2006 | AY597011.2 |
| 48 | | X | | Beta | Rousettus bat coronavirus HKU9-1 | China | 2006 | EF065513.1 |

| | | | | | | | | |
|---|---|---|---|---|---|---|---|---|
| 49 | | X | | Beta | Bat Hp-Betacoronavirus/Zhejiang2013 | China | 2013 | KF636752.1 |
| 50 | | X | | Beta | Rat-Betacoronavirus HKU24 | China | 2012 | KM349742.1 |
| 51 | | X | | Beta | Rousettus bat coronavirus | China | 2014 | KU762338.1 |
| 52 | | X | | Beta | Betacoronavirus Erinaceus / VMC / DEU | Germany | 2012 | KC545383.1 |
| 53 | | X | | Beta | Murine hepatitis virus A59 | Switzerland | 2004 | AY700211.1 |
| 54 | | X | | Beta | Human coronavirus OC43 | USA | 2004 | AY585228.1 |
| 55 | | X | | Beta | Murine hepatitis virus | USA | 1997 | AF029248.1 |
| 56 | | X | | Beta | Bovine coronavirus | USA | 2001 | AF391541.1 |
| 57 | | X | | Beta | Rat coronavirus Parker | USA | 2009 | FJ938068.1 |
| 58 | | X | | Beta | Rabbit coronavirus HKU-14 | China | 2006 | JN874559.1 |
| 59 | | X | | Beta | Betacoronavirus Erinaceus | Germany | 2012 | KC545386.1 |
| 60 | | X | | Beta | Homo sapiens Betacoronavirus England 1 | United Kingdom | 2012 | KC164505.2 |
| 61 | | X | | Beta | Homo sapiens Wuhan Hu-1 | Wuhan - China | 2019 | MN908947.1 |
| 62 | | X | | Beta | Homo sapiens SARS-CoV-2 | Wuhan - China | 2020 | MN938384.1 |
| 63 | | X | | Beta | Human coronavirus OC43 | USA | 2004 | AY585228 |
| 64 | | X | | Beta | Pipistrellus bat coronavirus HKU5 | China | 2006 | EF065509.1 |
| 65 | | X | | Beta | Tylonycteris bat coronavirus HKU4-1 | China | 2007 | EF065505.1 |
| 66 | | X | | Beta | Homo sapiens - pneumonia virus 2019nCoV/Japan | Japan | 2020 | LC521925.1 |
| 67 | | X | | Beta | Human betacoronavirus 2c EMC | Middle East | 2012 | JX869059.2 |
| 68 | | X | | Beta | Bat SARS HKU3-1 | China | 2003 | DQ022305 |
| 69 | | X | | Beta | Homo sapiens - pneumonia virus isolate WIV02 | Wuhan - China | 2019 | MN996527.1 |
| 70 | | X | | Beta | Homo sapiens- pneumonia virus isolate WIV04 | Wuhan - China | 2019 | MN996528.1 |
| 71 | | X | | Beta | Homo sapiens - pneumonia virus isolate WIV05 | Wuhan - China | 2019 | MN996529.1 |
| 72 | | X | | Alpha | Bat coronavirus CDPHE15/USA/2006 | USA | 2006 | KF430219.1 |

* Reference sequence of SARS-CoV-2 virus isolated from human at Wuhan, China.

**Table S2: Temporal exploration of 40 and 10 whole genome sequences using TempEST**

| **Function: Correlation** | **40 -Sequences** | | **10 Sequences** | |
|---|---|---|---|---|
| | *Arbitrary root* | *Best-fitting root* | *Arbitrary root* | *Best-fitting root* |
| *Slope* | -0.001 | 0.011 | 12.944 | 162.520 |
| *X - intercept* | 2286.71 | 1968.99 | 1780.31 | 1989.11 |
| *p-value* | 0.445 | 0.451 | 0.237 | 0.691 |
| $R^2$ | 0.198 | 0.203 | 0.056 | 0.477 |
| *Residual Mean* | 1.49E-04 | 2.02E-02 | 1.40E-05 | 1.45E-06 |

**Table S3: Characteristics of 20 core genomic fragments of length $L_f \geq 200$ obtained from 10 WGS. Number and percentage of Single Nucleotide Polymorphism (SNP's). Non-structural proteins (NSP's) identified from the 20 fragments along with their positional mapping to reference sequence of SARS-CoV-2 NC_045512.2 is presented.**

| Fragment Number | Fragment Length | No of Positions in Fragment with SNP's | Percentage of Positions in Fragment with SNP's | Total no of bases in Fragment across 10 WGS | Total no of SNP's in Fragment across 10 WGS | Percentage of SNP's in Fragment across 10 WGS | NSP's Positions in 10 WGS | NSP's in 10 WGS | Reference Sequence SARS-CoV- 2 NC_045512.2 NSP Positions |
|---|---|---|---|---|---|---|---|---|---|
| 1 | 393 | 142 | 36.13 | 3930 | 456 | 12.89 | 7388..7780 | Nsp3 | 2720..8554 |
| 2 | 236 | 78 | 33.05 | 2360 | 265 | 12.48 | 8305..8540 | Nsp3 | 2720..8554 |
| 3 | 246 | 99 | 40.24 | 2460 | 316 | 14.27 | 9183..9428 | Nsp4 | 8555..10054 |
| 4 | 220 | 83 | 37.73 | 2200 | 299 | 15.10 | 9540..9759 | Nsp4 | 8555..10054 |
| 5 | 208 | 66 | 31.73 | 2080 | 229 | 12.23 | 9783..9990 | Nsp4 | 8555..10054 |
| 6 | 247 | 72 | 29.15 | 2470 | 244 | 10.98 | 9991..10237 | Nsp4 & nsp5 | |
| | | | | | | | *9991..10054* | *Nsp4* | 8555..10054 |
| | | | | | | | *10055..10237* | *Nsp5* | 10055..10972 |
| 7 | 218 | 58 | 26.61 | 2180 | 223 | 11.37 | 11379..11596 | Nsp6 | 10973..11842 |
| 8 | 282 | 75 | 26.60 | 2820 | 274 | 10.80 | 11597..11878 | Nsp6 & nsp7 | |
| | | | | | | | *11597..11842* | *Nsp6* | 10973..11842 |
| | | | | | | | *11843..11878* | *Nsp7* | 11843..12091 |
| 9 | 482 | 107 | 22.20 | 4820 | 367 | 8.46 | 12171..12652 | Nsp8 | 12092..12685 |
| 10 | 701 | 153 | 21.83 | 7010 | 525 | 8.32 | 12788..13488 | Nsp9 &nsp10 &nsp12 | |
| | | | | | | | *12788..13024* | *Nsp9* | 12686..13024 |
| | | | | | | | *13025..13441* | *Nsp10* | 13025..13441 |
| | | | | | | | *13442..13468* | *Nsp12* | 13442..13468 |
| | | | | | | | *13468..13488* | *Join nsp12* | 13468..16236 |
| 11 | 658 | 192 | 29.18 | 6580 | 619 | 10.45 | 13563..14220 | Join nsp12 | 13468..16236 |
| 12 | 438 | 121 | 27.63 | 4380 | 374 | 9.49 | 14221..14658 | Join nsp12 | 13468..16236 |
| 13 | 560 | 131 | 23.39 | 5600 | 351 | 6.96 | 14681..15240 | Join nsp12 | 13468..16236 |
| 14 | 1611 | 382 | 23.71 | 16110 | 1244 | 8.58 | 15242..16852 | Join nsp12 & nsp13 | |
| | | | | | | | *15242..16236* | *Join nsp12* | 13468..16236 |
| | | | | | | | *16237..16852* | *Nsp13* | 16237..18039 |
| 15 | 1195 | 300 | 25.10 | 11950 | 982 | 9.13 | 16853..18047 | Nsp13 & nsp14 | |
| | | | | | | | *16853..18039* | *Nsp13* | 16237..18039 |
| | | | | | | | *18040..18047* | *Nsp14* | 18040..19620 |
| 16 | 202 | 68 | 33.66 | 2020 | 233 | 12.82 | 18387..18588 | Nsp14 | 18040..19620 |
| 17 | 650 | 193 | 29.69 | 6500 | 680 | 11.62 | 18732..19381 | Nsp14 | 18040..19620 |
| 18 | 233 | 78 | 33.48 | 2330 | 274 | 13.07 | 19890..20122 | Nsp15 | 19621..20658 |
| 19 | 479 | 131 | 27.35 | 4790 | 424 | 9.84 | 20430..20908 | Nsp15 & nsp16 | |
| | | | | | | | *20430..20658* | *Nsp15* | 19621..20658 |
| | | | | | | | *20659..20908* | *Nsp16* | 20659..20908 |
| 20 | 235 | 76 | 32.34 | 2350 | 237 | 11.21 | 21078..21312 | Nsp16 | 20659..21552 |

**Table S4: 4a, 4b and 4c - Propagation of Errors for 1st tMRCA, 2nd tMRCA and clock rate**

Supplementary Table S4a: Propagation of Errors for 1st tMRCA

| Evolutionary models | 1st tMRCA (t) | Node ages - height median | UCI | LCI | UCI-LCI= Delta(t) | $Delta(t)^2$ | | |
|---|---|---|---|---|---|---|---|---|
| 10WGS - CEP_RC | t1 | 13.77 | 20.34 | 8.87 | 11.47 | 131.56 | | |
| 20FG- CCP_RC | t2 | 10.59 | 17.52 | 7.39 | 10.13 | 102.62 | | |
| 20FG- CEP_RC | t3 | 9.68 | 15.13 | 7.48 | 7.65 | 58.52 | | |
| 8,11,12 FG - CCP_RC | t4 | 10.21 | 16.02 | 7.46 | 8.56 | 73.27 | | |
| 1,5,16FG - CCP_SC | t5 | 14.87 | 19.62 | 10.61 | 9.01 | 81.18 | | |
| 8,11,12 FG - CCP_SC | t6 | 21.9 | 28.29 | 16.03 | 12.26 | 150.31 | | |
| | | 13.50 | | | | 597.46 | 24.44 | **4.07** |
| | | | | | | Sum | Sqrt(Sum) | Sqrt(Sum)/6 |
| | | **13.5± 4.1** | | | | | | |

Supplementary Table S4b: Propagation of Errors for 2nd tMRCA

| Evolutionary models | 2nd tMRCA (t) | Node ages - height median | UCI | LCI | UCI-LCI= Delta(t) | $Delta(t)^2$ | | |
|---|---|---|---|---|---|---|---|---|
| 10WGS - CEP_RC | t1 | 22.32 | 33.24 | 12.61 | 20.63 | 425.60 | | |
| 20FG- CCP_RC | t2 | 17.07 | 35.29 | 9.16 | 26.13 | 682.78 | | |
| 20FG- CEP_RC | t3 | 15.29 | 24.36 | 9.18 | 15.18 | 230.43 | | |
| 8,11,12 FG - CCP_RC | t4 | 13.72 | 24.66 | 7.73 | 16.93 | 286.62 | | |
| 1,5,16FG - CCP_SC | t5 | 35.59 | 47.47 | 24.49 | 22.98 | 528.08 | | |
| 8,11,12 FG - CCP_SC | t6 | 48.28 | 61.05 | 34.8 | 26.25 | 689.06 | | |
| | | 25.38 | | | | 2842.57 | 53.32 | **8.89** |
| | | | | | | Sum | Sqrt(Sum) | Sqrt(Sum) / 6 |
| | | **25.4± 8.9** | | | | | | |

Supplementary Table S4c: Propagation of Errors for Clock rate

| Evolutionary models | Clock rate | UCI | LCI | UCI - LCI = Delta(t) | $Delta(t)^2$ | | |
|---|---|---|---|---|---|---|---|
| 10WGS - CEP_RC | 0.0029 | 0.0049 | 0.0018 | 0.00309 | 0.00000955 | | |
| 20FG- CCP_RC | 0.0039 | 0.0079 | 0.0022 | 0.0057 | 0.00003249 | | |
| 20FG- CEP_RC | 0.0059 | 0.0095 | 0.0041 | 0.0054 | 0.00002916 | | |
| 8,11,12 FG - CCP_RC | 0.0058 | 0.0116 | 0.0032 | 0.0084 | 0.00007056 | | |
| 1,5,16FG - CCP_SC | 0.0023 | 0.003 | 0.002 | 0.001 | 0.00000100 | | |
| 8,11,12 FG - CCP_SC | 0.0011 | 0.0014 | 0.001 | 0.0004 | 0.00000016 | | |
| | **0.00365** | | | | 0.00014292 | 0.01195484 | **0.001992473** |
| | | | | | Sum | Sqrt(Sum) | Sqrt(Sum)/6 |
| | **$3.65 \times 10^{-3} \pm 1.99 \times 10^{-3}$** | | | | | | |

**Table S5: Bayesian analysis clock rate estimates**

Bayesian clock rates obtained in section 4.3.2 for the 6 best fitting models selected by Nested sampling are estimated by the specification of clock rate estimates in Table S5. These were used to specify the uniform prior distribution on the clock rate using mean and 95% CI to conduct host probability analysis to estimate the clock rates results of Table 2 using BEAST 1.10.4

| Evolutionary models | Molecular clock rate |
|---|---|
| 10WGS - CEP_RC | 0.0074 [0.0018, 0.0116] |
| 20FG- CCP_RC | 0.0075 [0.0022, 0.0142] |
| 20FG- CEP_RC | 0.0179 [0.0041, 0.0374] |
| 8,11,12 FG - CCP_RC | 0.0126 [0.0032, 0.0246] |
| 1,5,16FG - CCP_SC | 0.0032 [0.0020, 0.0044] |
| 8,11,12 FG - CCP_SC | 0.0016 [0.0010, 0.0023] |

#### Supplementary Figures

(a)

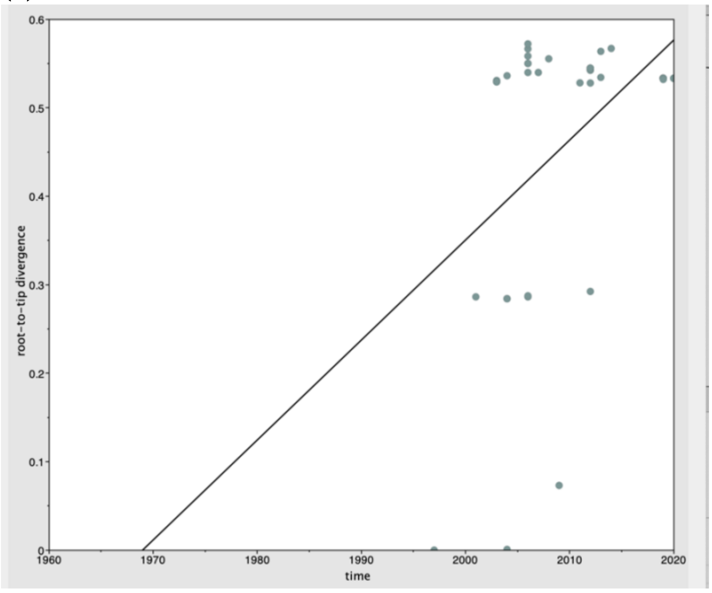

*40 Sequences – Best fitting root*
*($R^2$=0.203, Correlation Coefficient = 0.451)*

(b)

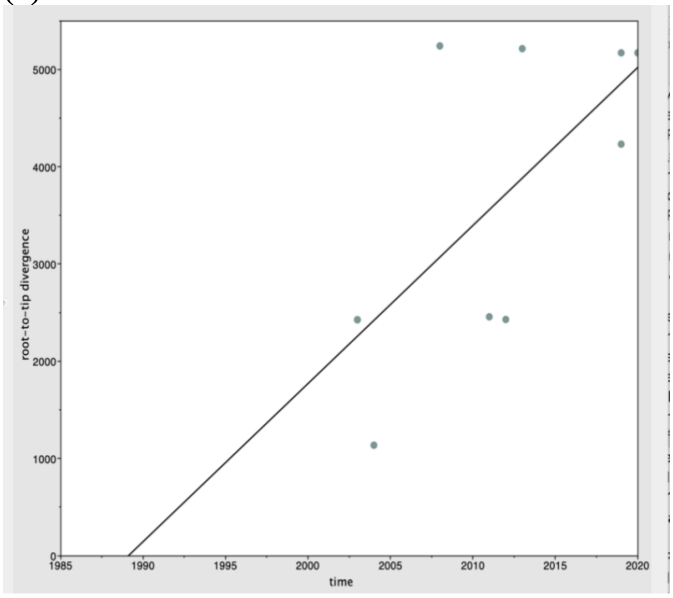

*10 Sequences – Best fitting root*
*($R^2$=0.477, Correlation Coefficient = 0.691)*

Supplementary Figure 1: Temporal Exploration of Sequences and Trees [TempEST] performed on Whole Genome Sequences

(a) 40 WGS and (b) 10 WGS. Root-to-tip divergence was estimated using TempEST. Correlation coefficient = 0.45 for 40 WGS and 0.691 for 10 WGS, which is not significant and indicates lack of temporal signal in both datasets.

(a)

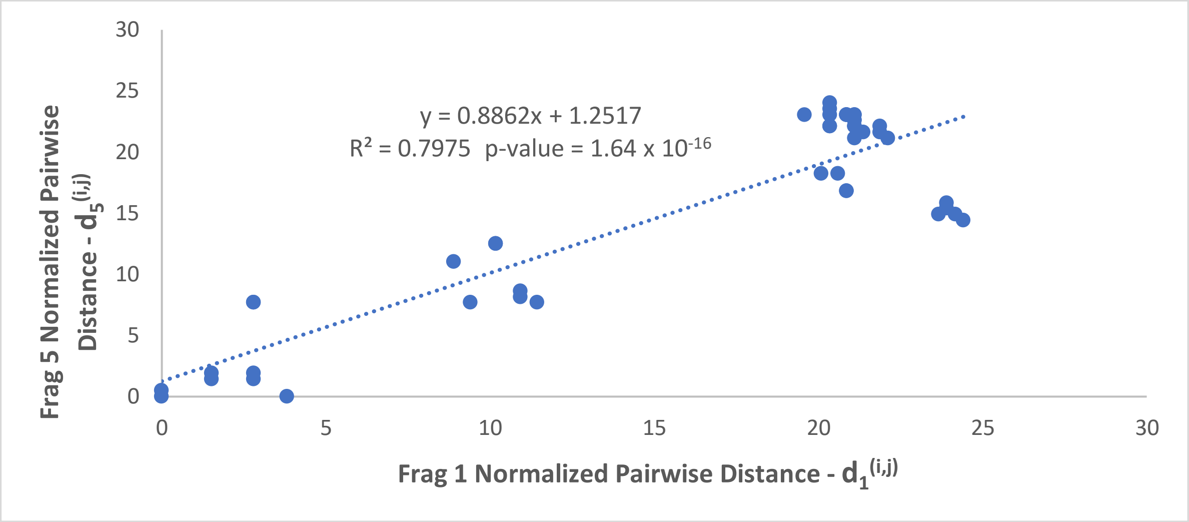

(b)

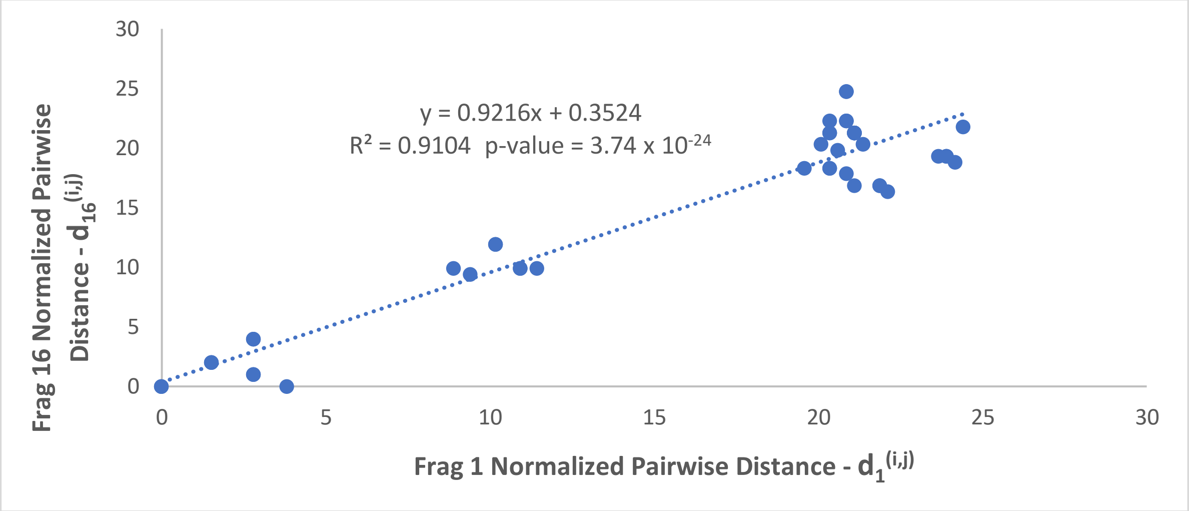

(c)

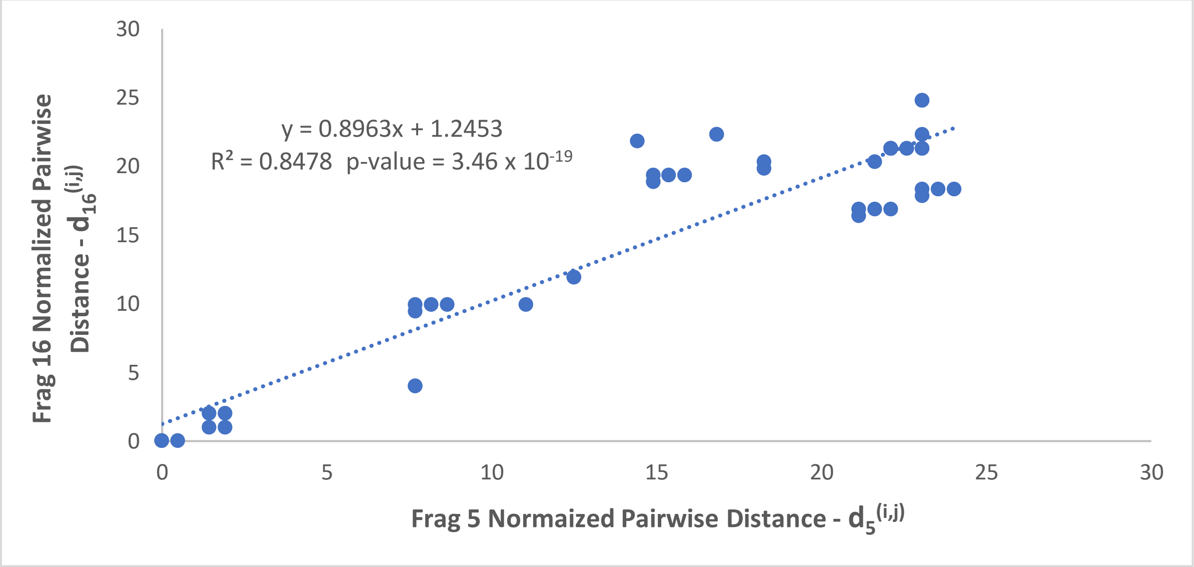

Supplementary Figure 2: Regression analysis of the normalized pairwise distance for fragment group {1,5,16}

This yielded a significant fit according to the criteria defined in section 3.2.2.2.2 of the main paper. The coordinates $(d_A^{(i,j)}, d_B^{(i,j)})$ of the points show the normalized pairwise distance between pairs of samples $(i, j)$ according to the fragments $A$ and $B$ indicated in the horizontal and vertical axis, respectively (see Figure 2(c)). (a) Fragment 1 against Fragment 5, (b) Fragment 1 against Fragment 16 and (c) Fragment 5 against Fragment 16. The dotted line shows to the fitted regression line.

(a)

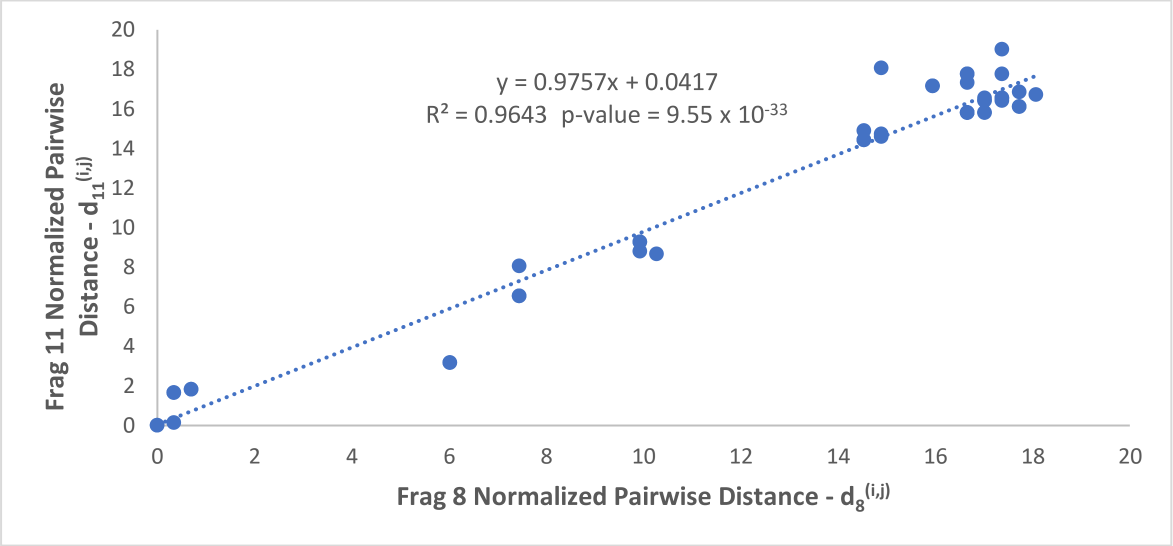

(b)

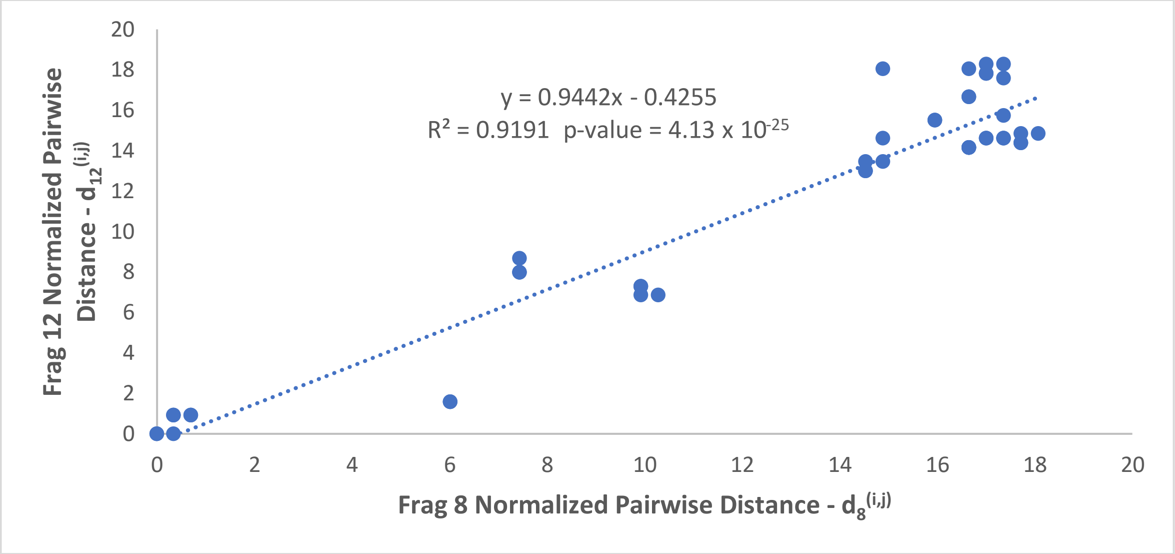

(c)

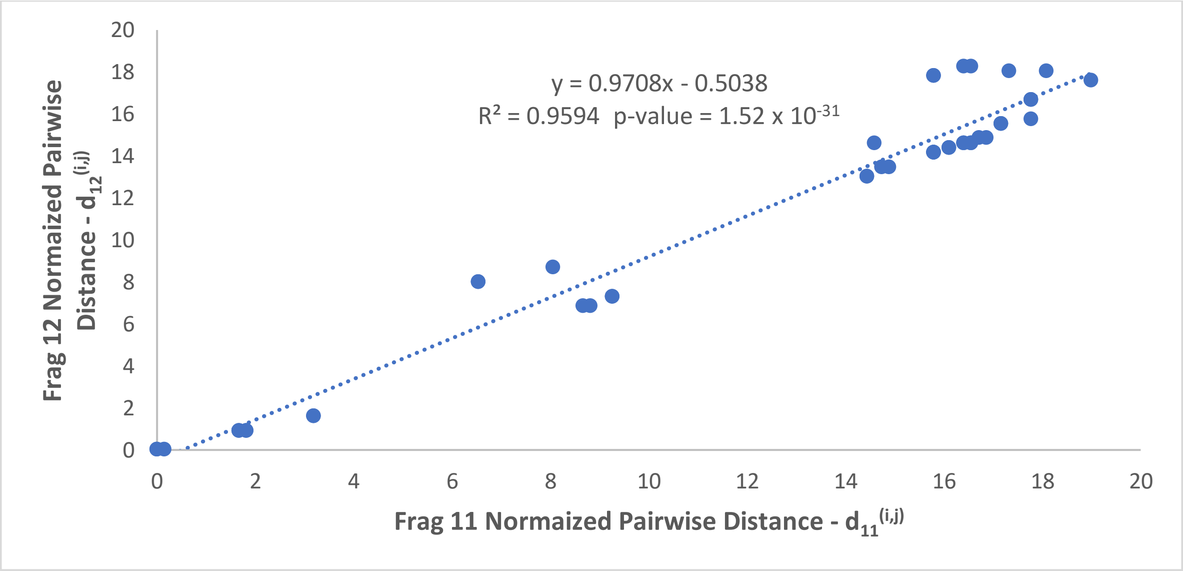

Supplementary Figure 3: Regression analysis of the normalized pairwise distance for fragment group {8, 11, 12}

This yielded a significant fit according to the criteria defined in section 3.2.2.2.2 of the main paper. The coordinates $(d_A^{(i,j)}, d_B^{(i,j)})$ of the points show the normalized pairwise distance between pairs of samples $(i, j)$ according to the fragments $A$ and $B$ indicated in the horizontal and vertical axis, respectively (see Figure 2 (c)). (a) Fragment 8 against Fragment 11, (b) Fragment 8 against Fragment 12 and (c) Fragment 11 against Fragment 12. The dotted line shows to the fitted regression line.

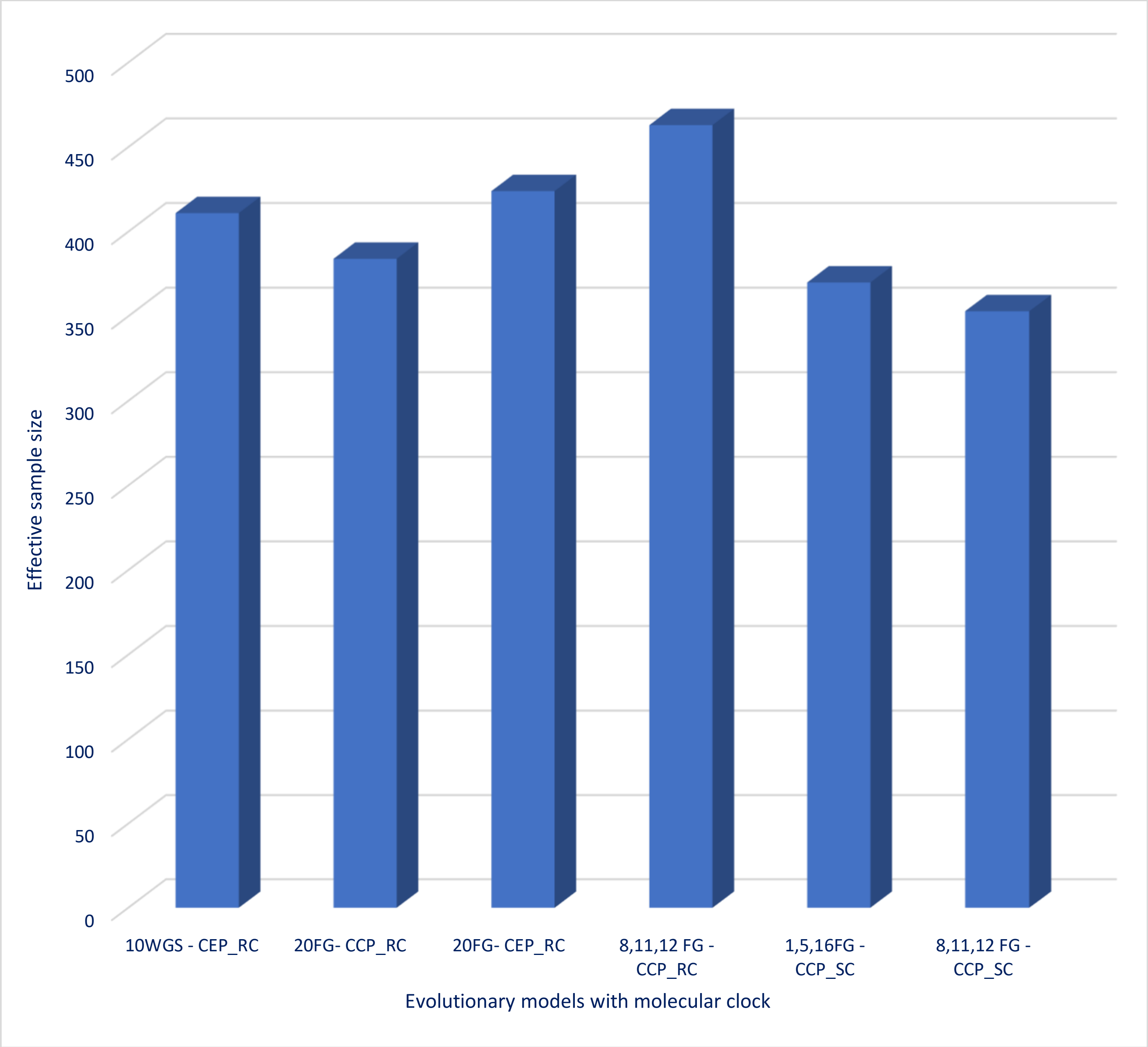

Supplementary Figure 4: BEAST Model selection by Nested Sampling - Estimation of Max effective sample size (ESS)

Results of Bayesian hypothesis testing for Bayesian evolutionary model selection conducted by Nested Sampling and comparison of Max ESS between strict and relaxed molecular clock for the final 6 best-fit models of 10 WGS, 20 fragments and 2 fragment groups.

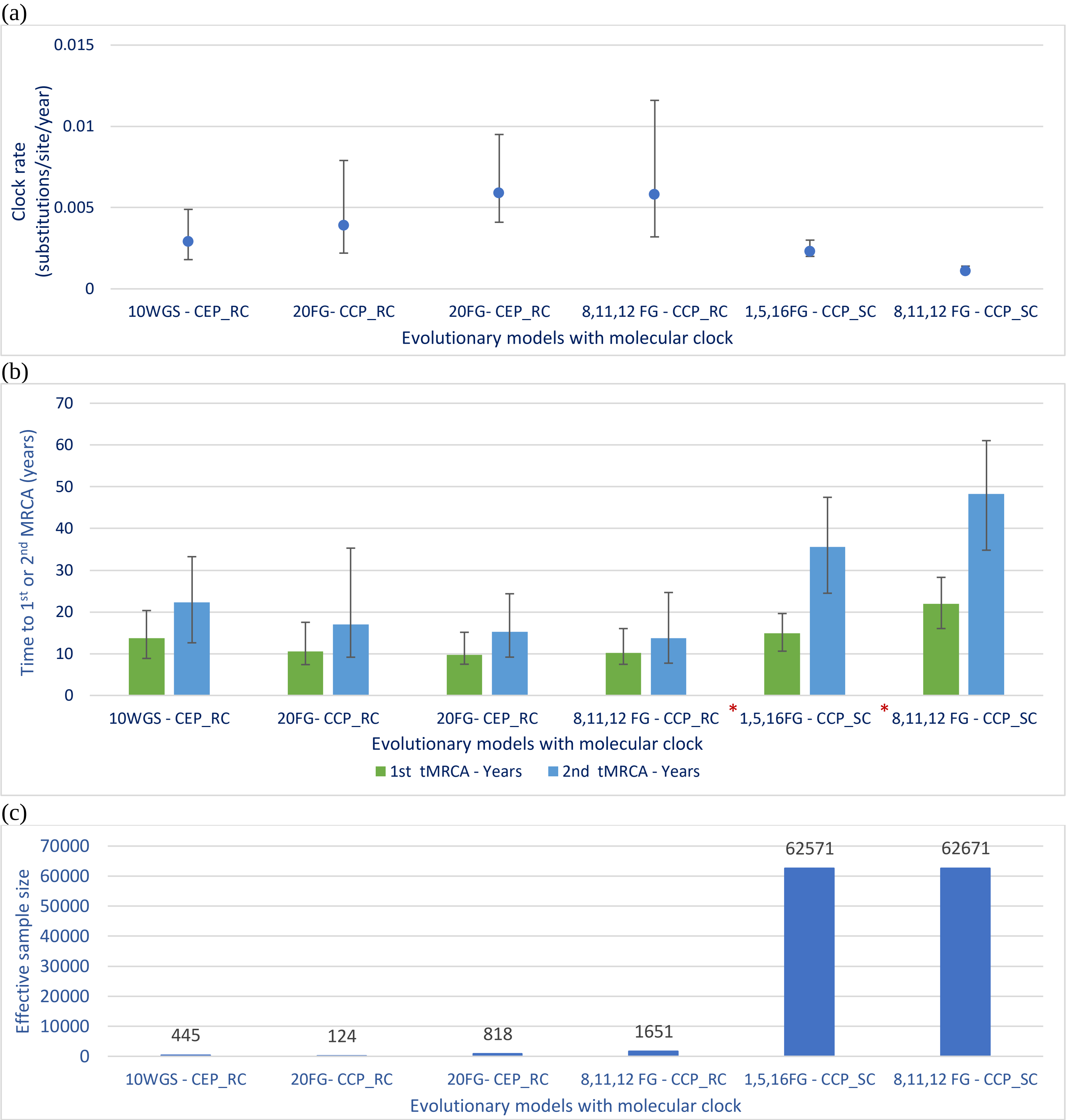

Supplementary Figure 5: BEAST Phylodynamic evolutionary analysis for the evolutionary models selected by Nested sampling

(a) Clock rate - Clock rate estimates for the mean clock rate with 95% highest posterior density (95% HPD) interval indicates that higher clock rates are obtained for 4 relaxed clock models when compared to the clock rates obtained for 2 strict clock models. (b) Evolutionary years to most recent common ancestor of SARS-CoV-2 - Time measured phylogenetic relationship of SARS-CoV-2 with closely related bat (green) and pangolin (blue) sequences: Estimated evolutionary years to most recent common ancestor of SARS-CoV-2 is the closest between SARS-CoV-2 and bat sequence RaTG13, Pangolin CoV is 2nd closest in time to SARS-CoV-2. Statistical significance is seen in strict molecular clock for divergence time estimates of Coalescent constant population evolutionary model for both fragment groups {1,5,16} and {8,11,12}, as the confidence intervals of 1st tMRCA do not overlap with those of 2nd tMRCA. (c) Effective sample size - Results of estimation for Whole genome sequences versus fragment groups comparison with strict molecular clock and relaxed molecular clock. See Table 2: Results of the Bayesian phylodynamic analyses for the best fitting models identified by nested sampling.

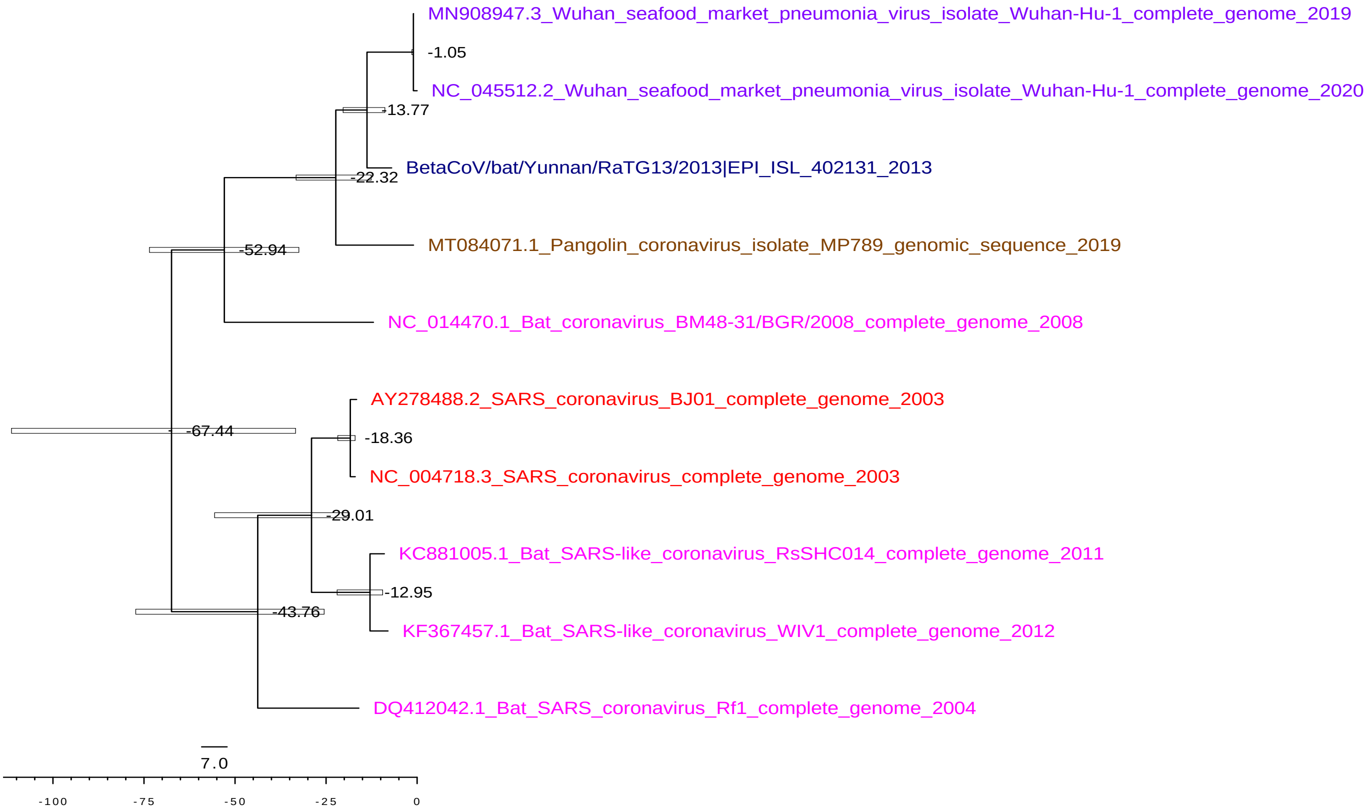

Supplementary Figure 6.1: 10WGS CEP RC (scale is in years)
BEAST Phylotraits Temporal Phylogeny for 10WGS with Coalescent Exponential Population and HKY sub-models and relaxed molecular clock. Bat coronavirus RaTG13 (indigo) (-13.77 years) is the 1st tMRCA of SARS-CoV-2 and Pangolin (brown) (-22.32 years) is the 2nd tMRCA to SARS-CoV-2 (purple), SARS-CoV (red), other beta CoV (pink). The blue bars are indicating 95% credible interval for all internal node ages. (Dataset: Table S1, Selection 3)

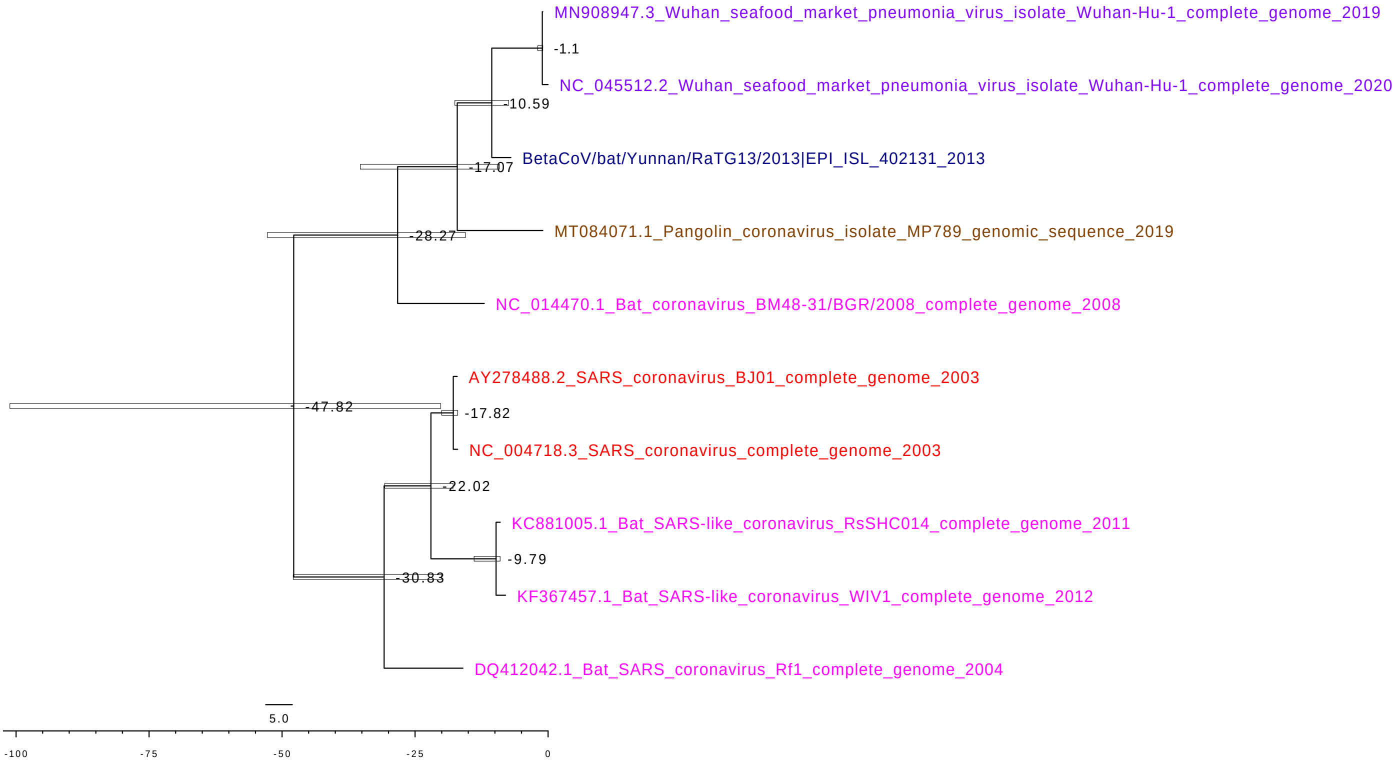

Supplementary Figure 6.2: 20FG_CCP_RC (scale is in years)
BEAST Phylotraits Temporal Phylogeny for the 20 fragments group with Coalescent Constant Population and HKY sub-models and relaxed molecular clock. Bat coronavirus RATG13(indigo) (-10.59 years) is the 1st tMRCA of SARS-CoV-2 and Pangolin (brown) (-17.07 years) is the 2nd tMRCA to SARS-CoV-2 (purple), SARS-CoV (red), other beta CoV (pink). The blue bars are indicating 95% credible interval for all internal node ages. (Dataset: Table S1, Selection 3)

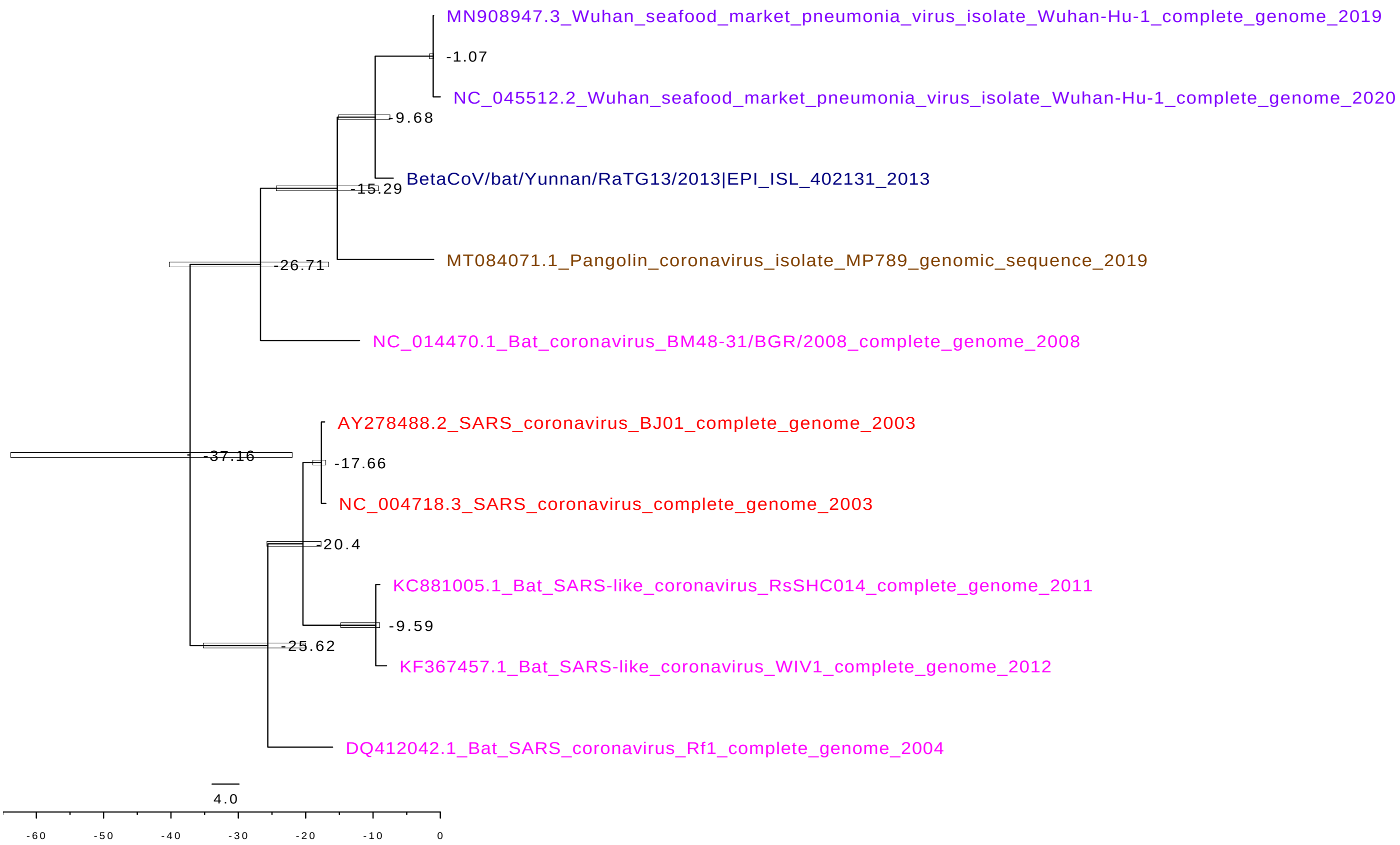

Supplementary Figure 6.3: 20FG_CEP_RC (scale is in years)
BEAST Phylotraits Temporal Phylogeny for the 20 fragments group with Coalescent Exponential Population and HKY sub-models and relaxed molecular clock. Bat coronavirus RATG13 (indigo) (-9.68 years) is the 1st tMRCA of SARS-CoV-2 and Pangolin (brown) (-15.29 years) is the 2nd tMRCA to SARS-CoV-2 (purple), SARS-CoV (red), other beta CoV (pink). The blue bars are indicating 95% credible interval for all internal node ages. (Dataset: Table S1, Selection 3)

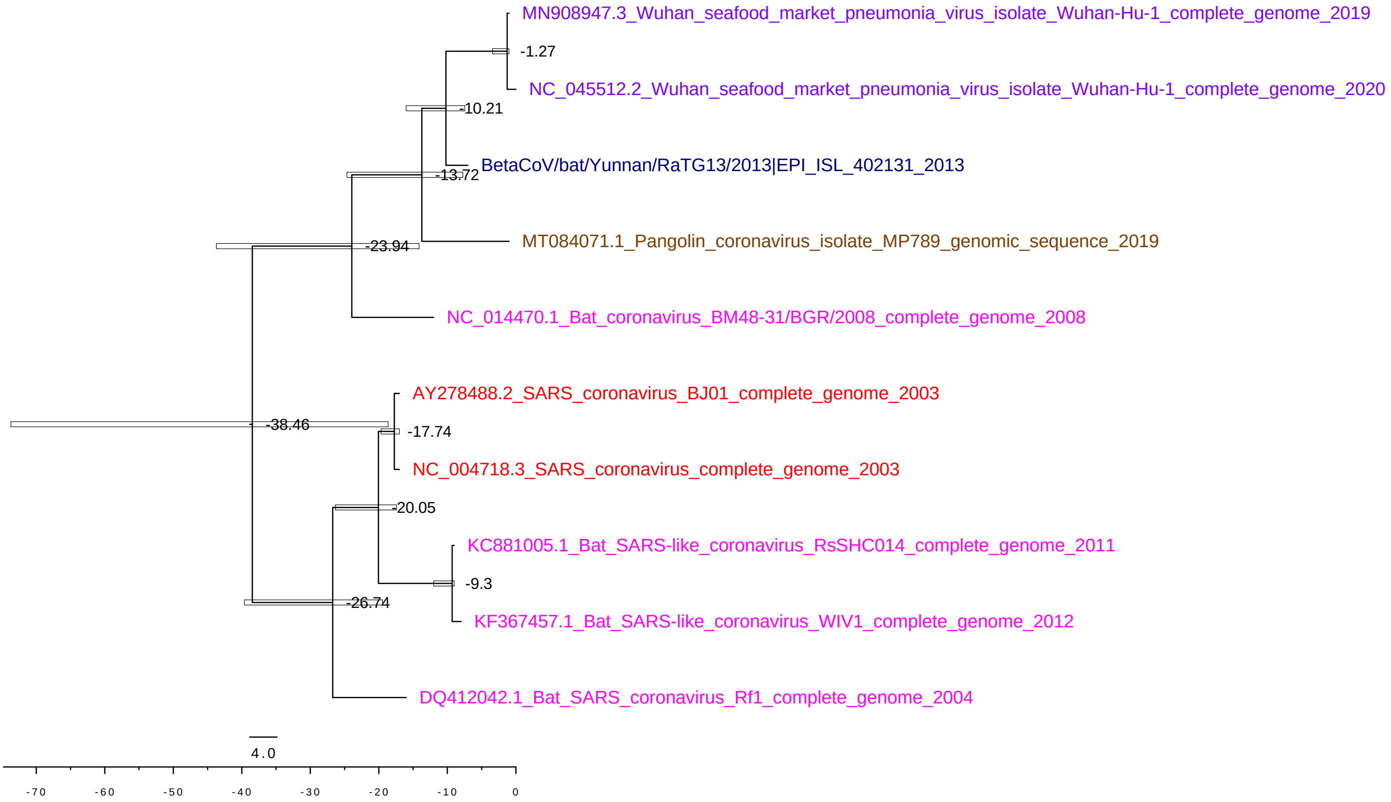

Supplementary Figure 6.4: 8,11,12 CCP_RC (scale is in years)
BEAST Phylotraits Temporal Phylogeny for the fragment group {8,11,12} with a Coalescent Constant Population and HKY sub-models and relaxed molecular clock. Bat coronavirus RATG13(indigo) (-10.21 years) is the 1st tMRCA of SARS-CoV-2 and Pangolin (brown) (-13.72 years) is the 2nd tMRCA to SARS-CoV-2 (purple), SARS-CoV (red), other beta CoV (pink). The blue bars are indicating 95% credible interval for all internal node ages. (Dataset: Table S1, Selection 3)

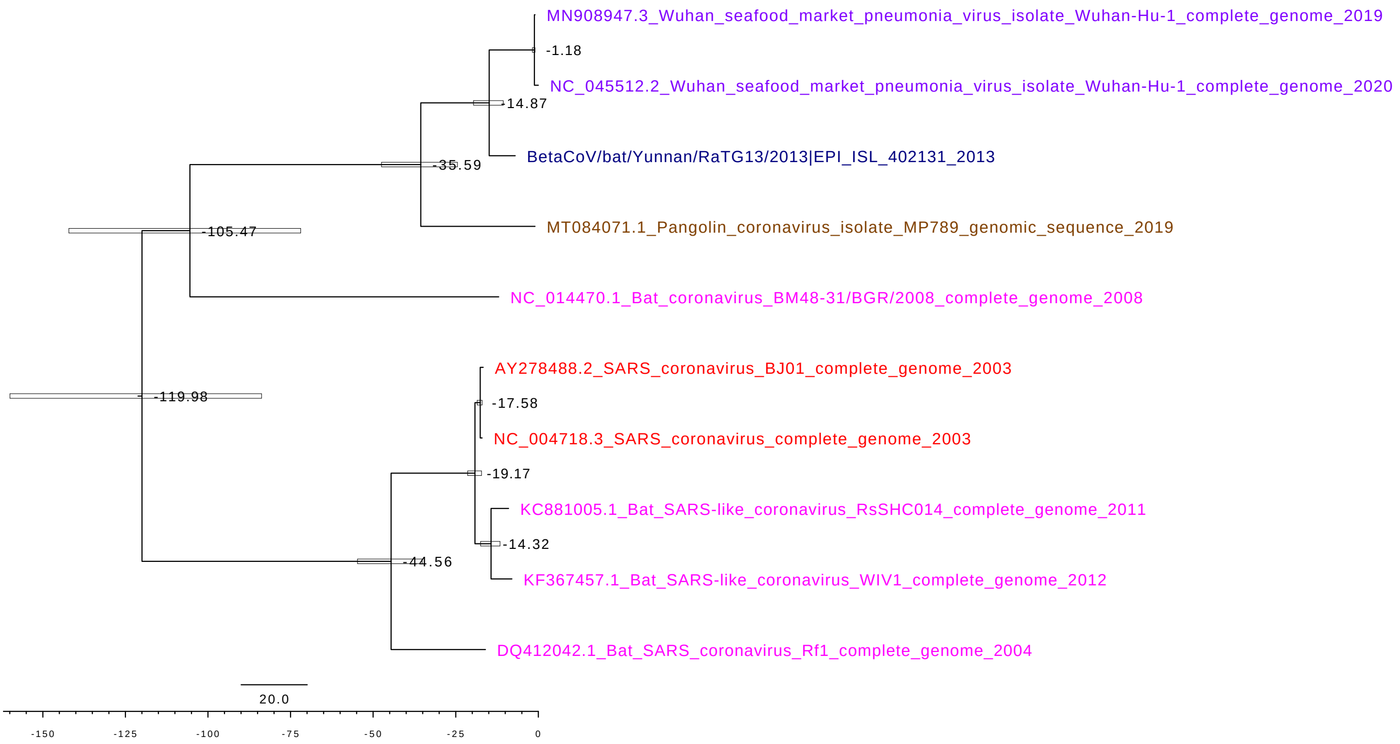

Supplementary Figure 6.5: 1,5,16FG_CCP_SC (scale is in years)
BEAST Phylotraits Temporal Phylogeny for the fragment group {1,5,16} with Coalescent constant Population and HKY sub-models and strict molecular clock. Bat coronavirus RATG13(indigo) (-14.87 years) is the 1st tMRCA of SARS-CoV-2 and Pangolin (brown) (-35.59 years) is the 2nd tMRCA to SARS-CoV-2 (purple), SARS-CoV (red), other beta CoV (pink). The blue bars are indicating 95% credible interval for all internal node ages. (Dataset: Table S1, Selection 3)

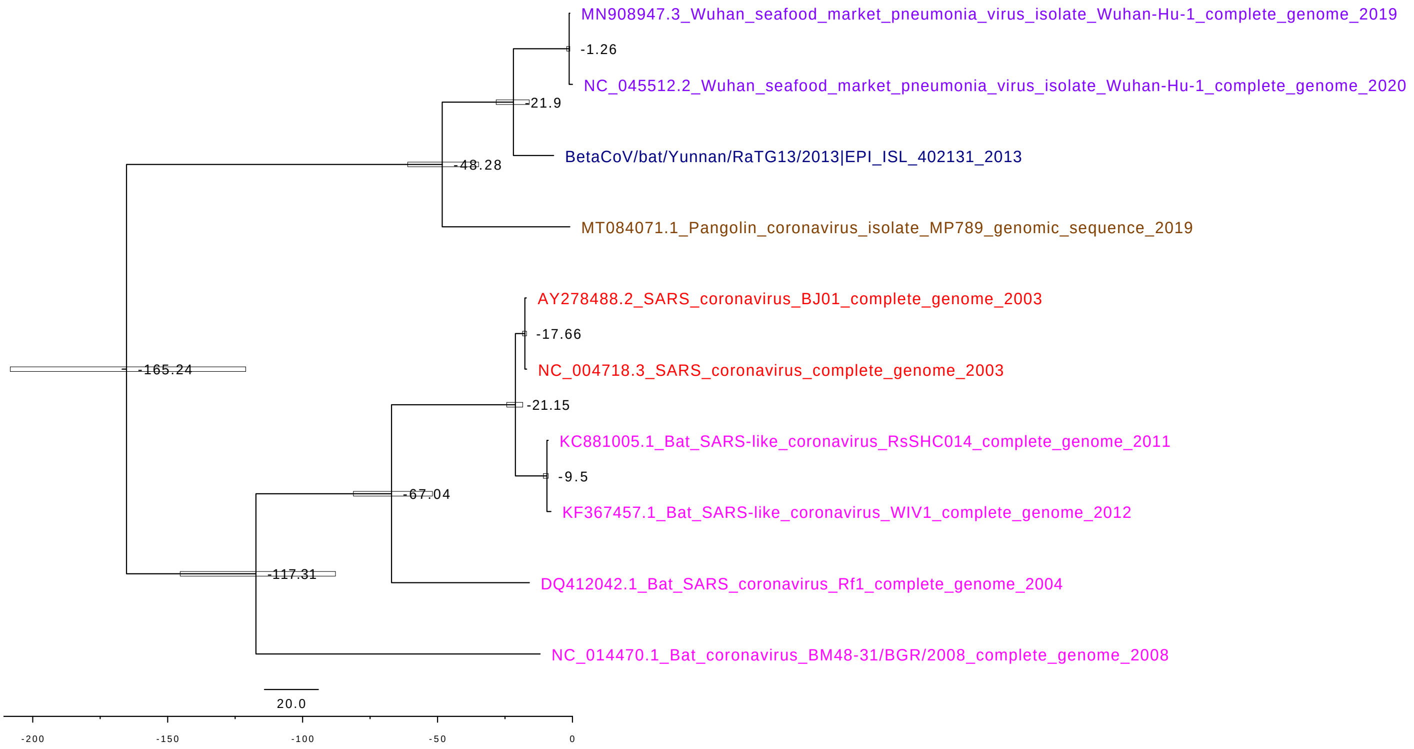

Supplementary Figure 6.6: 8,11,12FG_CCP_SC (scale is in years)
BEAST Phylotraits Temporal Phylogeny for the fragment group {8,11,12} with Coalescent Constant Population and HKY sub-models and strict molecular clock. Bat coronavirus RATG13 (indigo) (-21.9 years) is the 1st tMRCA of SARS-CoV-2 and Pangolin (brown) (-48.28 years) is the 2nd tMRCA to SARS-CoV-2 (purple), SARS-CoV (red), other beta CoV (pink). The blue bars are indicating 95% credible interval for all internal node ages. (Dataset: Table S1, Selection 3)

Supplementary Figure 6: BEAST Phylotraits Evolutionary Analysis maximum clade credibility phylodynamic trees

(a)

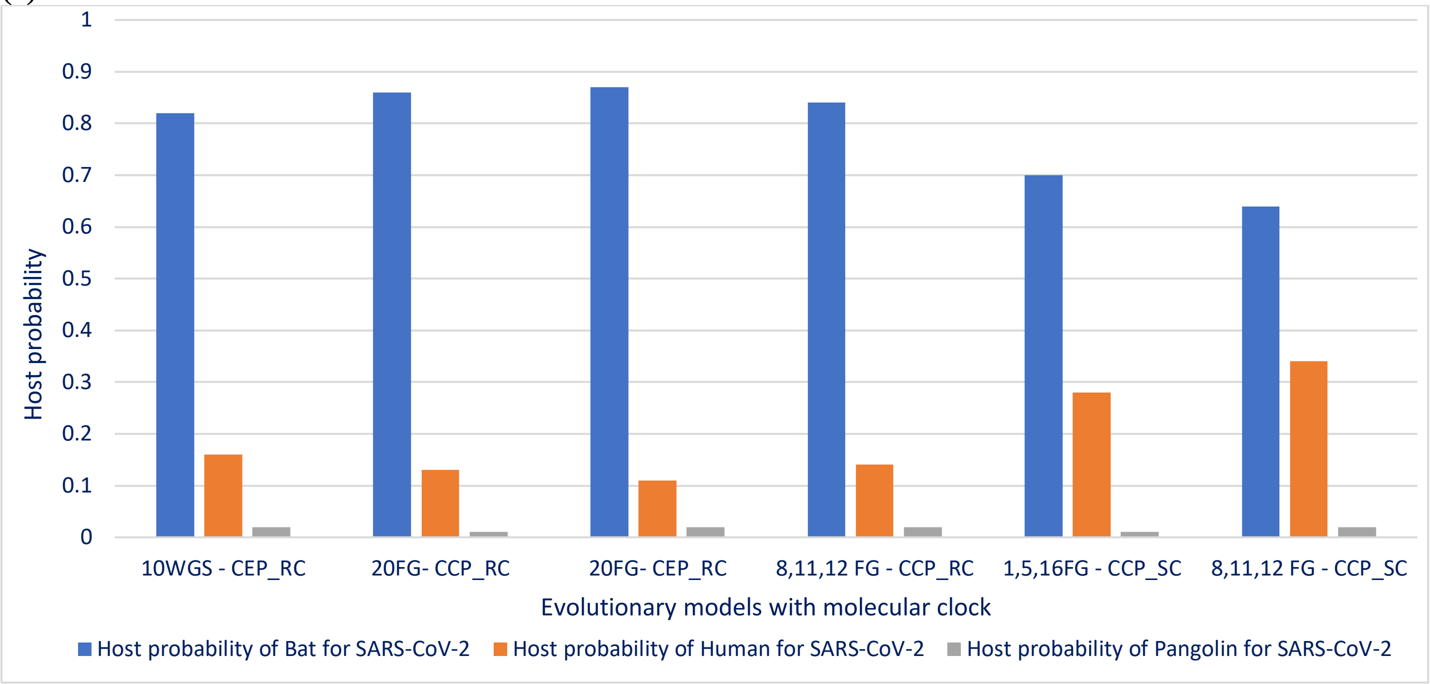

(b)

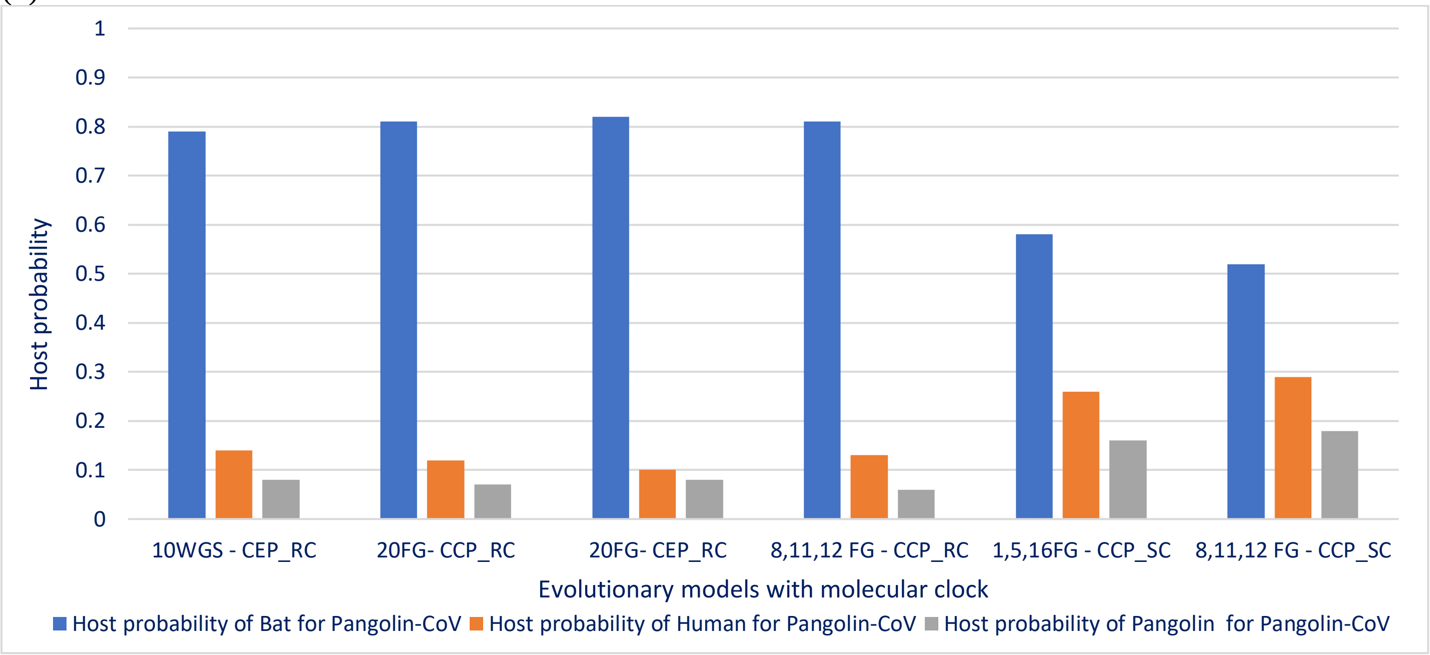

Supplementary Figure 7: BEAST Phylotraits evolutionary analysis – Host probability for SARS-CoV-2 and Pangolin CoV

(a) Host probability for SARS-CoV-2 - Host probability phylogenetic diffusion process suggests that SARS-CoV-2 emerged in natural zoonotic origin of bats (blue) identified as the most likely reservoir host of SARS-CoV-2

(b) Host probability for Pangolin-CoV - Host probability suggests that Pangolin-CoV (gray) also emerged from zoonotic origin of bats, as bat was identified again as the most likely reservoir host of Pangolin

See Table 2: Results of the Bayesian phylodynamic analyses for the best fitting models identified by nested sampling.

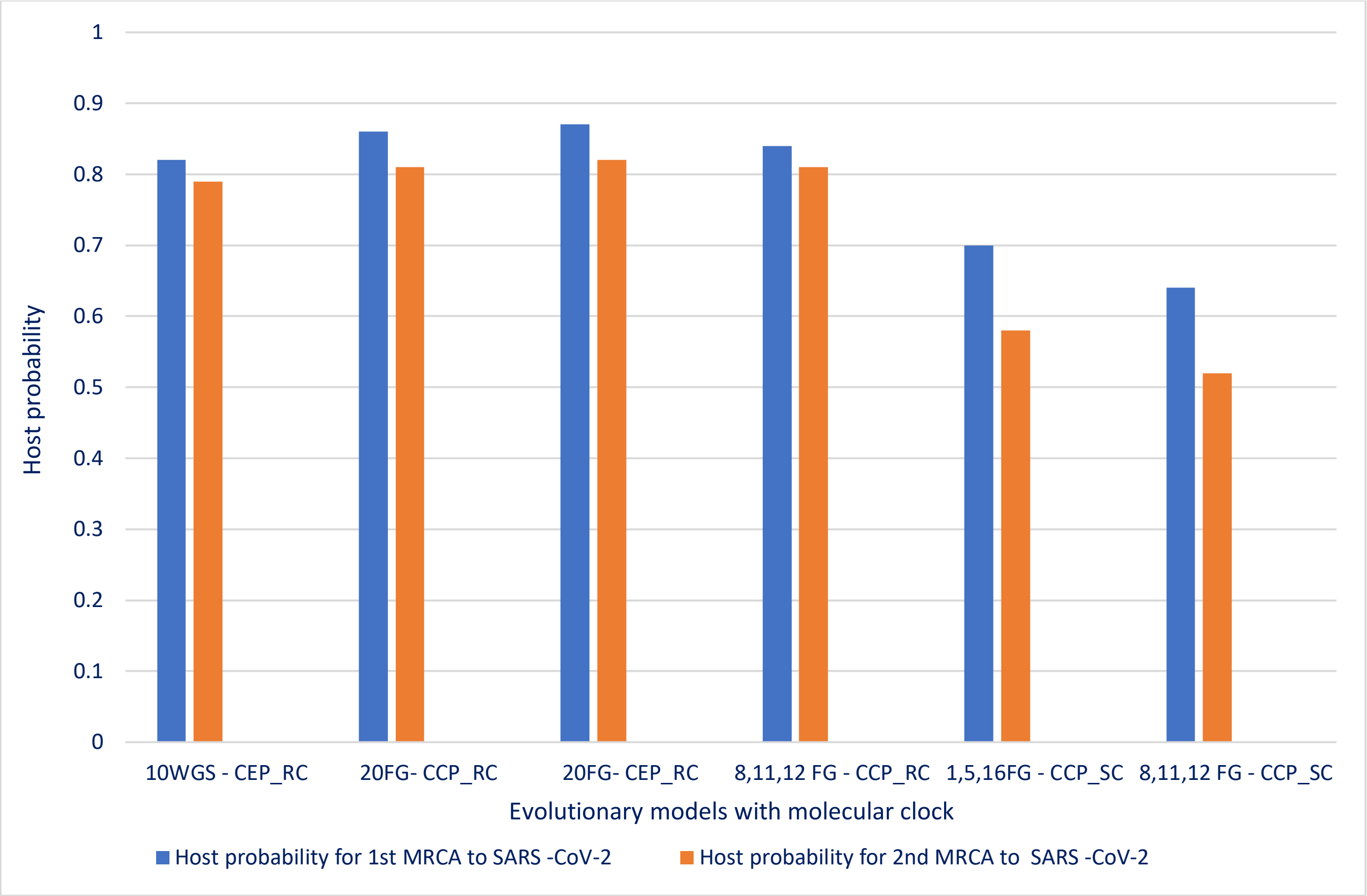

Supplementary Figure 8: BEAST Phylotraits evolutionary analysis Host Probability of MRCA to SARS-CoV-2

Blue – Host probability of Bat as 1$^{st}$ MRCA for SARS-CoV2
Orange - Host probability of Pangolin as 2$^{nd}$ MRCA for SARS-CoV2

The results indicate that SARS-CoV-2 probably emerged from bats, as it was identified as the most likely 1$^{st}$ MRCA for all six models. Pangolin was 2$^{nd}$ MRCA for SARS-CoV2 for all six models.
